# NMR Resonance Assignment of the LA Motif of Human LA-Related Protein 1

**DOI:** 10.1101/2023.05.09.539749

**Authors:** Benjamin C. Smith, Robert Silvers

## Abstract

Human La-related protein 1 (HsLARP1) is involved in post-transcriptional regulation of certain 5’
s terminal oligopyrimidine (5’TOP) mRNAs as well as other mRNAs and binds to both the 5’TOP motif and the 3’-poly(A) tail of certain mRNAs. HsLARP1 is heavily involved in cell proliferation, cell cycle defects, and cancer, where HsLARP1 is significantly upregulated in malignant cells and tissues. Like all LARPs, HsLARP1 contains a folded RNA binding domain, the La motif (LaM). Our current understanding of post-transcriptional regulation that emanates from the intricate molecular framework of HsLARP1 is currently limited to small snapshots, obfuscating our understanding of the full picture on HsLARP1 functionality in post-transcriptional events. Here, we present the nearly complete resonance assignment of the LaM of HsLARP1, providing a significant platform for future NMR spectroscopic studies.

## Introduction

La-related proteins (LARPs) constitute a highly conserved superfamily of RNA-binding proteins that are essential in controlling the destiny of many RNAs. The significance of this particular family of proteins is that close to all known eukaryotic organisms have retained members of the LARP superfamily in their genomes.[1] Of the five distinct classes of LARPs identified within the human genome, LARPs 1, 4, and 6 are predominantly located in the cytosol and are involved in post-transcriptional gene control, thus regulating the delicate balance between active translation, degradation, and storage of mRNAs.[1-12] The intricate nature of post-transcriptional gene control responsible for maintaining the delicate balance between active translation, degradation, and storage of mRNAs makes it a target for dysfunction on many levels. Hence, many LARPs are intimately implicated in a variety of diseases including cancer, fibrosis, and neurodegenerative diseases.[3, 4, 6, 7, 13-18]

Phylogenetic studies indicate that the majority of members of the LARP superfamily including HsLARP1, contain a folded RNA binding domain, the La motif (LaM), that together with a downstream RNA-recognition motif (RRM) connected by a short linker form a region called the La-module[1, 19]. Human La-related protein 1 (HsLARP1) is involved in post-transcriptional regulation of certain 5’
s terminal oligopyrimidine (5’TOP) mRNAs as well as other mRNAs,[3-5] and biochemical studies found that the La-module of HsLARP1 binds to both the 5’TOP motif and the 3’-poly(A) tail of certain mRNAs.[20, 21] It was shown that HsLARP1 activity is regulated through the mammalian target of rapamycin (mTOR) pathway.[4, 22-24] Furthermore, several independent studies indicated that HsLARP1 is heavily involved in cell proliferation, cell cycle defects,[25, 26] and cancer, where HsLARP1 is significantly upregulated in malignant cells and tissues.[4, 13-16] HsLARP1 has been found to be upregulated in ovarian cancers and there is some evidence of its importance in cell motility which could be important for some metastatic cancer.[27, 28]

Interestingly, HsLARP1 was also found to be involved in the cellular response of various viral infections. HsLARP1 was shown to associate with the nucleocapsid of SARS-CoV-2[29] and also bind to SARS-CoV-2 genome and repress replication of SARS-CoV-2. [30] While the genome of SARS-CoV-2 lacks a 5’TOP motif, it contains some pyrimidine enriched regions in the 5’-untranslated region of its genome.[30] Furthermore, HsLARP1 has also been shown to be upregulated in Dengue virus infection[31] and has been found to associate with the surface of Hepatitis C virions.[32] HsLARP1 has also been found to interact with the capsid of Zika Virus.[33]

Our current understanding of post-transcriptional regulation that emanates from the intricate molecular framework of HsLARP1 is currently limited to small snapshots, obfuscating our understanding of the full picture on HsLARP1 functionality in post-transcriptional events. Here, we present the nearly complete resonance assignment of the LaM of HsLARP1, providing a significant platform for future NMR spectroscopic studies.

## Experimental Section

### Construct design

The HsLARP1(392-486) construct used in this study spans the residues S392 to P486 of the full-length isoform 1 of human La-related protein 1 (Uniprot Q6PKG0). HsLARP1(392-486) was commercially synthesized with a codon optimized sequence and inserted into a pET28a(+) vector (Gene Universal, USA) so that the construct initially produced recombinantly contains an N-terminal His_6_-tag followed by a tobacco etch virus (TEV) cleavage site. The TEV protease recognition sequence ENLYFQ▾S was chosen to retain the natural residue S392 after His_6_-tag removal.

### Expression of HsLARP1(392-486)

The plasmid coding for His_6_-TEV-HsLARP1(392-486) was transformed into chemically competent *E. coli* One Shot™ BL21 Star™ (DE3) cells (Invitrogen). Freshly transformed cells were grown on M9 agar containing 1 g/L ^1^NH_4_Cl, 10 g/L D-glucose, 1.5% agar, and 40 μg/mL kanamycin at 37°C overnight. 50 ml M9 minimal medium precultures were grown at 37°C up to an OD_600_ of 1.0 and used to inoculate main cultures. Uniformly [^13^C,^15^N]-labeled His_6_-TEV-HsLARP1(392-486) was expressed using M9 minimal medium containing 1 g/L ^15^NH_4_Cl, 2 g/L ^13^C-D-glucose, and 40 μg/mL kanamycin. Uniformly [^15^N-Leu]-, [^15^N-Phe]-, [^15^N-Ile]-, [^15^N-Val]-, and [^13^C,^15^N-Arg]-labeled His_6_-TEV-HsLARP1(392-486) were expressed using protocols published previously [34, 35]. Uniformly [^15^N, 2-^13^C-glycerol]-labeled and [^15^N, 1,3-^13^C-glycerol]-labeled His_6_-TEV-HsLARP1(392-486) were expressed using protocols published previously [36-38]. After inoculation, the main cultures were allowed to grow at 37°C in an orbital shaker set to 225 rpm until an OD_600_ of around 0.6 was reached and were then transferred to a water-ice bath for 15 min. After the cold shock, isopropyl-β-D-1-thiogalactopyranoside (IPTG) was added to a final concentration of 1 mM and the flasks were transferred to an orbital shaker set to 18°C and 225 rpm. Expression at 18°C was allowed to proceed for around 14 hours. Cells were harvested at 4°C and 5,000 × g for 30 min by centrifugation using a Beckman Coulter™ Avanti™ J-20 XPI centrifuge equipped with a JLA-8.1000 rotor. The supernatant was discarded and cell pellets were resuspended in a buffer containing 25 mM Tris/HCl pH 8.0, 400 mM NaCl, 0.1% w/v 3-[(3-cholamidopropyl)dimethylammonio]-1-propanesulfonate (CHAPS), 50 mM imidazole, 10% v/v glycerol. Protease inhibitors AEBSF, E-64, leupeptin, and aprotinin were added at final concentrations of 500 μM, 1 μM, 1 μM, and 150 nM, respectively. Cell suspensions were either stored at -20°C or subjected to purification immediately.

### Purification of HsLARP1(392-486)

Cells were lysed by running the cell suspension through a Microfluidizer® M-110L (Microfluidics) in 3-5 cycles at a pressure of 50 PSI while continuously cooling the lysate in a water ice bath. The lysate was cleared of cell debris by centrifugation at 4°C and 50,000 × g for 30 min by centrifugation using a Beckman Coulter™ Avanti™ J-20 XPI centrifuge equipped with a JA-25.50 rotor. Subsequently, the supernatant was passed through a 0.45 μm pore filter and applied at a flow rate of 5 ml/min to a 5 mL HisTrap™ HP column (Cytiva, USA) previously equilibrated with His-Trap buffer A containing 25 mM Tris/HCl pH 8.0, 0.1% w/v CHAPS, 400 mM NaCl, 50 mM Imidazole, and 10% v/v glycerol. The column was then washed at a flow rate of 5 ml/min with His-Trap buffer A until the UV-signal at 260 nm and 280 nm reached a stable baseline. The flowthrough was discarded. HsLARP1(392-486) was eluted by running His-Trap buffer B containing 25 mM Tris/HCl pH 8.0, 0.1% w/v CHAPS, 400 mM NaCl, 500 mM Imidazole, and 10% v/v glycerol through the column at a flow rate of 5 ml/min. During elution, 5 ml fractions were collected and fractions containing His6-TEV-HsLARP1(392-486) were pooled. His6-TEV-HsLARP1(392-486) was subjected to TEV digestion by adding 1 mL of 1 mg/mL TEV protease and placing it in a Spectra/Por® 3 dialysis membrane with a molecular weight cutoff (MWCO) of 3.5kDa. The sample was dialyzed against 1 L of a buffer containing 25 mM Tris/HCl pH 8.0, 150 mM NaCl, and 5 mM β-mercaptoethanol at 4°C overnight.

The His-tag, the His-tagged TEV protease and impurities were removed by reverse His-Trap column. The digested sample was applied at a flow rate of 5 ml/min to a 5 mL HisTrap™ HP column (Cytiva, USA) previously equilibrated with His-Trap buffer A. The flowthrough was collected. The column was subsequently washed with His-Trap buffer A until the UV-signal at 260 nm and 280 nm reached a stable baseline. Flowthrough and wash fractions containing HsLARP1(392-486) were pooled and mixed with an equal volume of Q-Buffer A containing 25 mM Tris/HCl pH 8.0, 2 mM EDTA, and 5 mM β-mercaptoethanol. The sample was then applied to a HiTrap™ Q FF (Cytiva, USA) column previously equilibrated with 90% Q-Buffer A and 10% Q-Buffer B containing 25 mM Tris/HCl pH 8.0, 1.0 M NaCl, 2 mM EDTA, and 5 mM β-mercaptoethanol. The flowthrough was collected. A wash with 90% Q-Buffer A and 10% Q-Buffer B was performed and the wash was collected. While the isoelectric point of HsLARP1(392-486) is predicted to be around 5.2 (ExPASy server ProtParam tool - http://web.expasy.org/protparam) and thus expected to bind to the Q column under the chosen conditions, it did not bind to the Q column and was found in the flowthrough. Flowthrough and wash fractions were combined and concentrated using a Vivaspin® 20 3kDa MWCO spin concentrator (Cytiva, USA).

Afterwards, the sample was subjected to size exclusion chromatography (SEC) using a HiLoad 26/600 Superdex 75 pg column (Cytiva, USA) previously equilibrated with SEC Buffer containing 22 mM Tris/HCl pH 7.5 and 110 mM NaCl. The sample was subsequently applied at a flowrate of 1.0 ml/min and fractions containing HsLARP1(392-486) were combined. Sample purity was assessed by SDS-PAGE. An extinction coefficient of ε_280_ = 11,460 M^-1^cm^-1^ was determined using the ExPASy server ProtParam tool (http://web.expasy.org/protparam) for HsLARP1(392-486) and employed for determining protein concentrations.

### NMR methods

Samples of HsLARP1(392-486) were prepared in a buffer containing 20 mM Tris/HCl pH 7.5, 100 mM NaCl, 10% D_2_O for NMR spectroscopic studies. 3-(Trimethylsilyl)-1-propanesulfonic acid sodium salt (DSS) was added to all samples for internal ^1^H referencing. ^13^C and ^15^N resonances were referenced indirectly[39]. NMR samples were prepared in either 5 mm thin-wall NMR tubes (Wilmad 541 PP-7) or 5mm Shigemi BMS-005B NMR tubes (Shigemi Co., Japan) at concentrations ranging from 230 μM to 1.3 mM. All NMR spectra were acquired at 298 K on a 700 MHz Bruker Advance III NMR spectrometer equipped with a TCI cryoprobe. Chemical shift assignments of backbone and sidechain resonances of HsLARP1(392-486) were conducted on uniformly [^13^C, ^15^N]-labelled samples, unless stated otherwise. A list of NMR experiments can be found in Table S1. For the backbone assignment, three-dimensional triple-resonance experiments such as HNCA[40-43], HNCACB[44, 45], HNCO[46-48], and HN(CA)CO[47, 49] were employed. Additionally, [^1^H, ^15^N]-HSQC spectra of uniformly [^15^N-Leu]-, [^15^N-Phe]-, [^15^N-Ile]-, [^15^N-Val]-, and [^13^C,^15^N-Arg]-labeled HsLARP1(392-486) samples were used as reference points for backbone assignment. For sidechain assignments, 3D (H)CC(CO)NH[50-55], 3D H(CCCO)NH[50-55], and 3D NOESY-HSQC spectra were employed. Additionally, to resolve overlapping resonances, [^1^H, ^13^C]-HSQC and 3D NOESY-HSQC spectra of uniformly [^15^N, 2-^13^C-glycerol]-labeled and [^15^N, 1,3- ^13^C-glycerol]-labeled were used. Data were processed with TopSpin 4.2.0 (Bruker BioSpin) and analyzed using Sparky (T. D. Goddard and D. G. Kneller, SPARKY 3, University of California, San Francisco). Secondary structure and torsional angles were predicted from chemical shifts using TALOS-N.[56]

## Results and Discussion

### NMR Resonance Assignment

The HsLARP1(392-486) construct used in this study contained a total of 95 residues including 4 prolines with a molecular weight of approximately 11.3 kDa. We used a combination of 3D experiments for backbone assignments, namely 3D HNCACB, HNCA, HNCO, and HN(CA)CO experiments. Amino-acid specific labeling ([^15^N-Leu], [^15^N-Phe], [^15^N-Ile], [^15^N-Val], and [^13^C,^15^N-Arg]) was used to provide resonance identity, thus aiding in unambiguous resonance assignment. The [^1^H-^15^N]-HSQC spectra of the amino acid-selectively labelled samples are shown in Figures S1 and S2. Of the expected 90 ^1^H-^15^N amide resonances in the [^1^H-^15^N]-HSQC spectrum (Figure 1), we were able to assign 88/90 resonances (∼97.8%). The backbone amide resonances of T393 and H445 are missing, most likely due to exchange broadening. Using the assignment strategy outlined above, a total of 98.7% of backbone resonances (88/90 H, 88/90 N, 93/95 C, 95/95 C^α^, 95/95 H^α^) were assigned.

**Figure 1:**
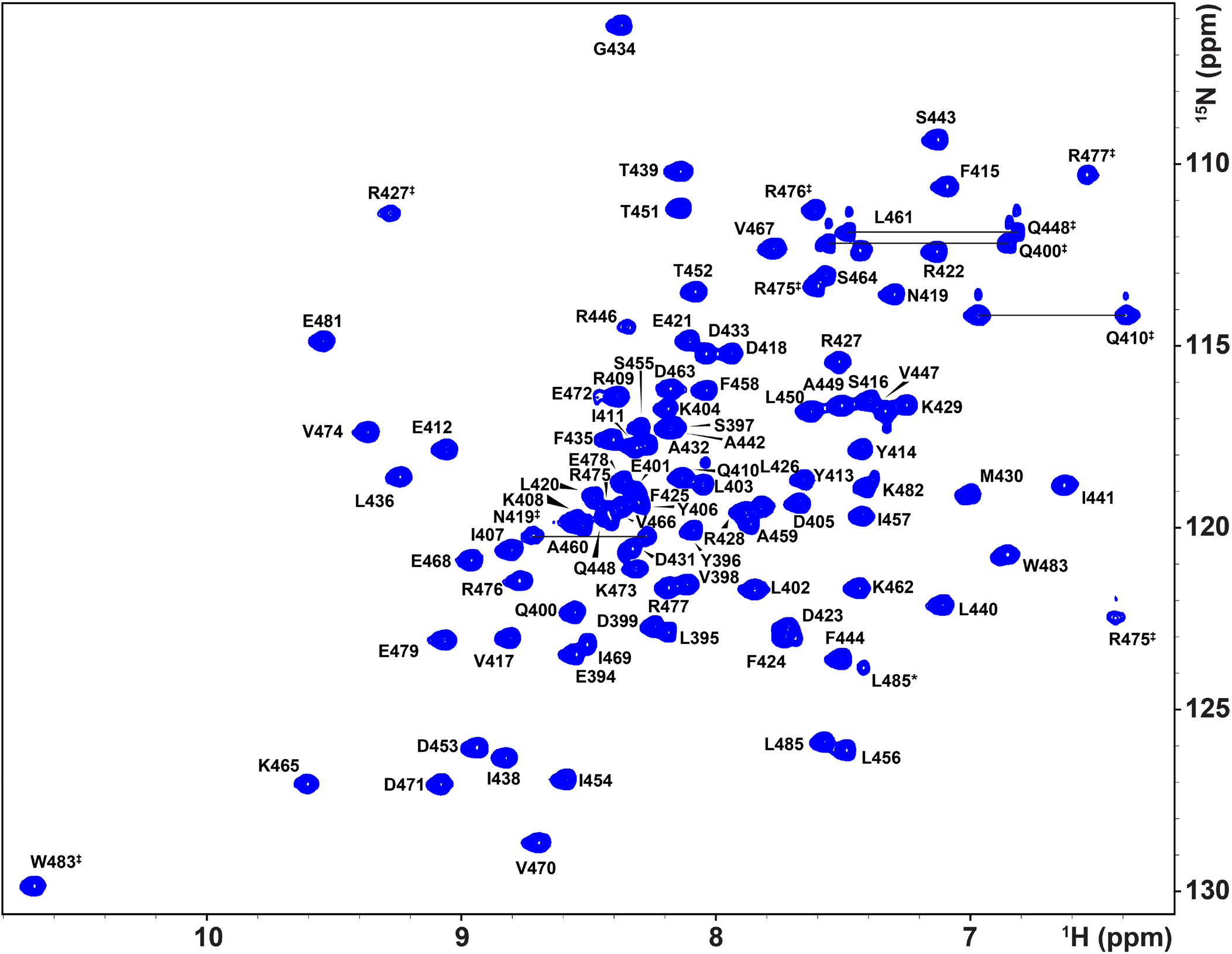
[^1^H,^15^N]- HSQC spectrum of 1.3 mM [^13^C,^15^N]-labeled HsLARP1(392-486) recorded at 700 MHz and 298 K in a buffer containing 20 mM Tris/HCl pH 7.5, 100 mM NaCl, 10% D_2_O. Side chain resonances are labeled with a double dagger (‡), peak splitting of L485 is indicated by an asterisk (*). Side chain resonances belonging to arginine are folded.

The side chain amide resonances in the [^1^H-^15^N]-HSQC spectrum (Figure 1) stemming from all asparagine (N419) and glutamine (Q400, Q410, and Q448) residues were assigned using a combination of 3D HNCACB and 3D NOESY-^15^N-HSQC spectra. Similarly, the ^1^H^ε1^-^15^N ^ε1^ resonance of W483 was assigned. The H^ε^-N^ε^ resonances of four of the eight arginine side chains are visible in the [^1^H-^15^N]-HSQC spectrum (Figure 1) and were assigned using a combination of 3D HNCACB and 3D NOESY-^15^N-HSQC spectra. Resonance assignment of aliphatic protons was achieved using a combination of 3D (H)CC(CO)NH and 3D H(CCCO)NH spectra as well as aliphatic 3D NOESY-^13^C-HSQC spectra of [^15^N, 2-^13^C-glycerol]-labeled and [^15^N, 1,3-^13^C-glycerol]-labeled HsLARP1(392-486). Using 1,3-^13^C glycerol or 2-^13^C glycerol as carbon source introduces ^13^C on alternating carbons for the majority of amino acids. This results in [^1^H-^13^C]-HSQC spectra with fewer resonances and more narrow lines (Figures S5 and S6). The assignment of side chain arginine resonance was aided by a [^1^H-^13^C]-HSQC spectrum of [^13^C,^15^N-Arg]-labeled HsLARP1(392-486) (Figure S3). The large number of aromatic residues (∼12%) in HsLARP1(392-486) leads to a significant overlap of aromatic peaks in [^1^H-^13^C]-HSQC spectra (Figure S4, black). Aromatic [^1^H-^13^C]-HSQC spectra and 3D NOESY-^13^C-HSQC spectra of [^15^N, 2-^13^C-glycerol]-labeled and [^15^N, 1,3-^13^C-glycerol]-labeled HsLARP1(392-486) were used to assign all aromatic protons. Using the assignment strategy outlined above, a total of 100% aliphatic and 100% aromatic resonances of non-exchangeable protons were assigned. Overall, 99.4% of protons in HsLARP1(392-486) were assigned. All chemical shifts of HsLARP1(392-486) are listed in Table S2.

### Chemical Shift Analysis

Secondary structure in HsLARP1(392-486) was predicted by TALOS-N[56] using the chemical shifts of H, H^α^, C, C^α^, C^β^, and N nuclei (Figure 2). TALOS-N predicts the secondary structure topology of the αααβαααββ fold known for the La motifs of other LARPs. However, an additional short helix is predicted for residues P480-K482, that is unique to HsLARP1(392-486). We used STRIDE[57] to determine the secondary structure of an AlphaFold prediction available from the AlphaFold Protein Structure Database[58, 59]. The secondary structure predicted by TALOS-N and AlphaFold are in good agreement. Interestingly, AlphaFold predicts the presence of a short 3_10_-helix in the same location as TALOS-N (P480-K482).

**Figure 2:**
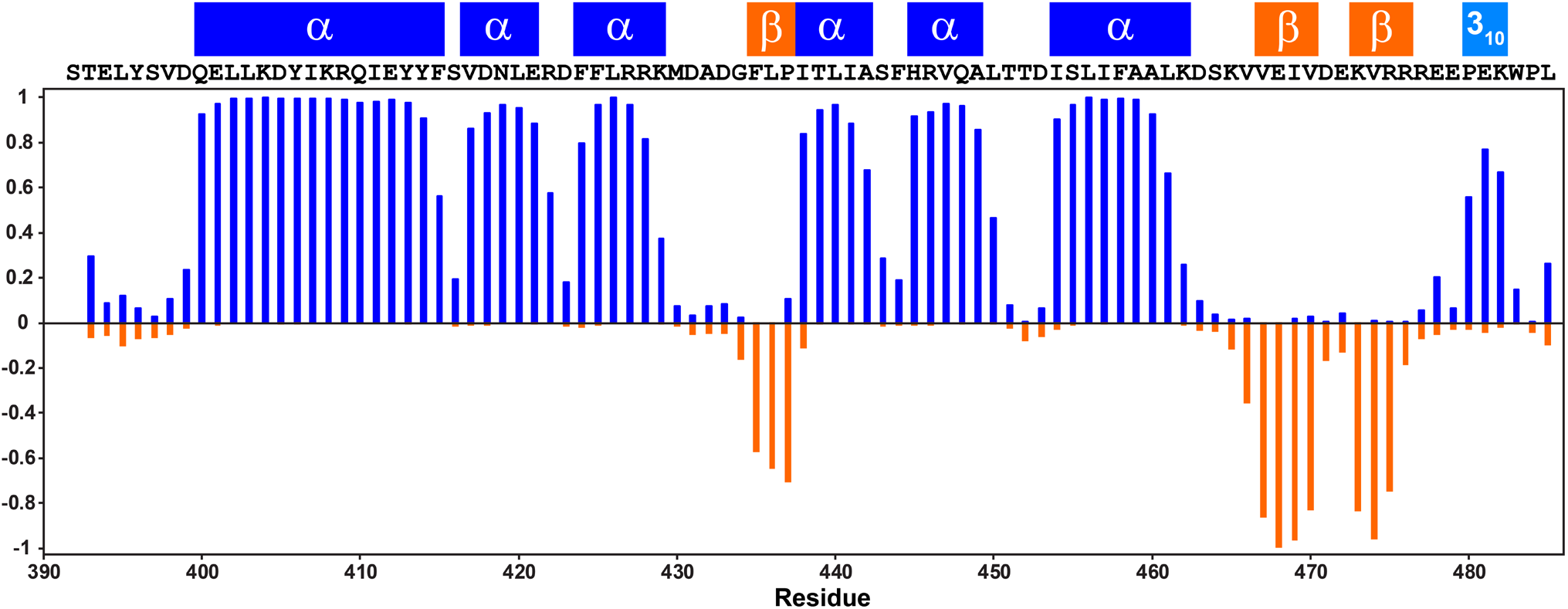
Secondary structure of HsLARP1(392-486). Chemical shifts were used to determine the α-helix propensity (shown in blue from 0 to 1) and β-strand propensity (shown in orange from 0 to -1). The amino acid sequence is shown. STRIDE was used to indicate location of α-helices, β-strands, and a 3_10_-helix in an AlphaFold prediction of HsLARP1(392-486).

## Acknowledgements

Research reported in this publication was supported by NIGMS of the National Institutes of Health under award number R35GM142912. The content is solely the responsibility of the authors and does not necessarily represent the official views of the National Institutes of Health.

