## Supporting Information for "NMR Resonance Assignment of the LA Motif of Human LA-Related Protein 1"

### Supporting Figures

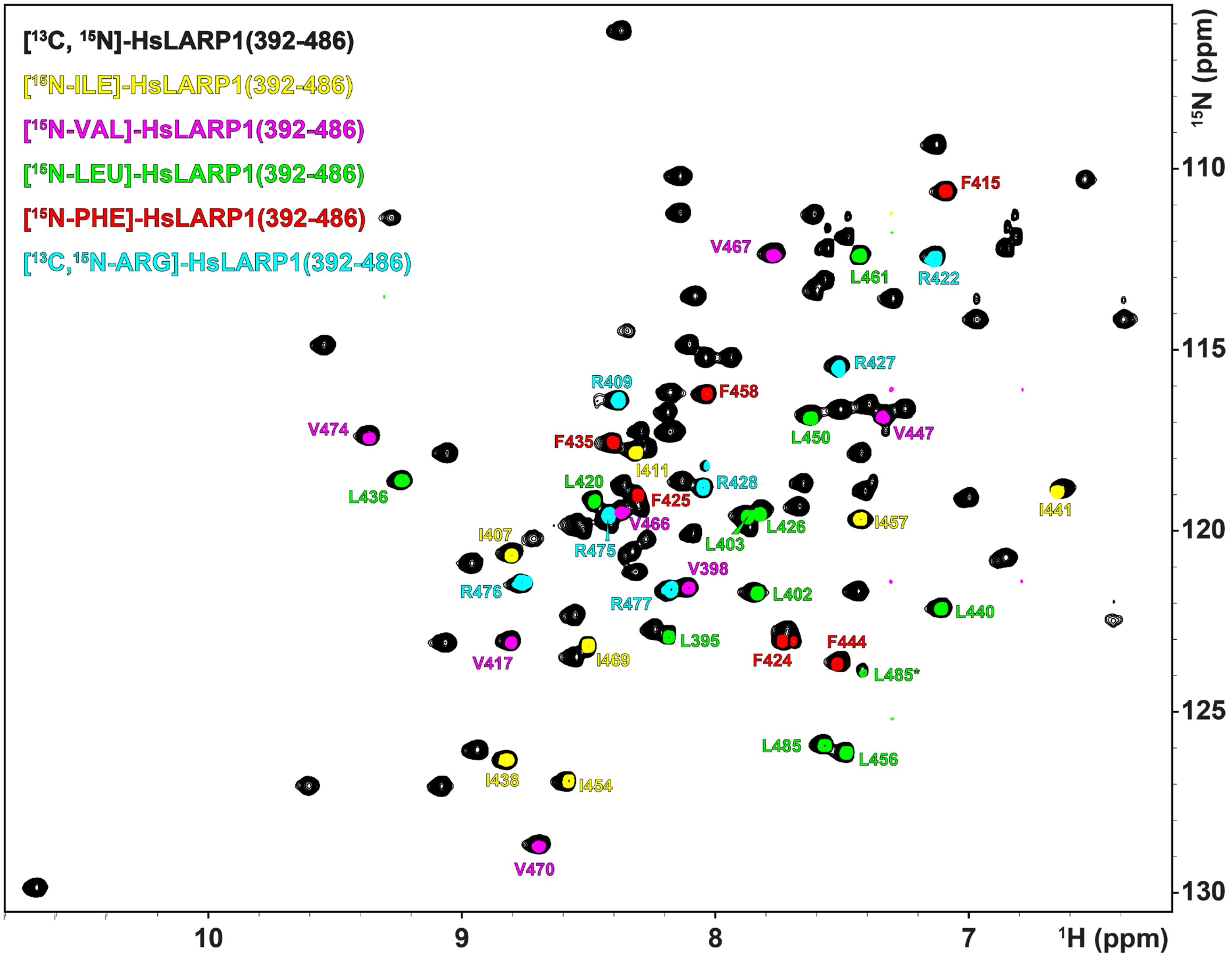


**Figure S1.** [$^1$H,$^{15}$N]-HSQC spectra of amino acid-specific labeling. The [$^1$H,$^{15}$N]-HSQC spectrum of 1.3 mM [$^{13}$C,$^{15}$N]-labeled HsLARP1(392-486) (black), the [$^1$H,$^{15}$N]-HSQC spectrum of 380 µM [$^{15}$N-Ile]-labeled HsLARP1(392-486) (yellow), the [$^1$H,$^{15}$N]-HSQC spectrum of 480 µM [$^{15}$N-Val]-labeled HsLARP1(392-486) (pink), the [$^1$H,$^{15}$N]-HSQC spectrum of 420 µM [$^{15}$N-Leu]-labeled HsLARP1(392-486) (green), the [$^1$H,$^{15}$N]-HSQC spectrum of 340 µM [$^{15}$N-Phe]-labeled HsLARP1(392-486) (yellow), and the [$^1$H,$^{15}$N]-HSQC spectrum of 230 µM [$^{13}$C,$^{15}$N-Arg]-labeled HsLARP1(392-486) (cyan) are shown. All spectra were recorded at 700 MHz and 298 K using a sample in a buffer containing 20 mM Tris/HCl pH 7.5, 100 mM NaCl, 10% $D_2O$. Amino acid-specifically labelled resonances are indicated.

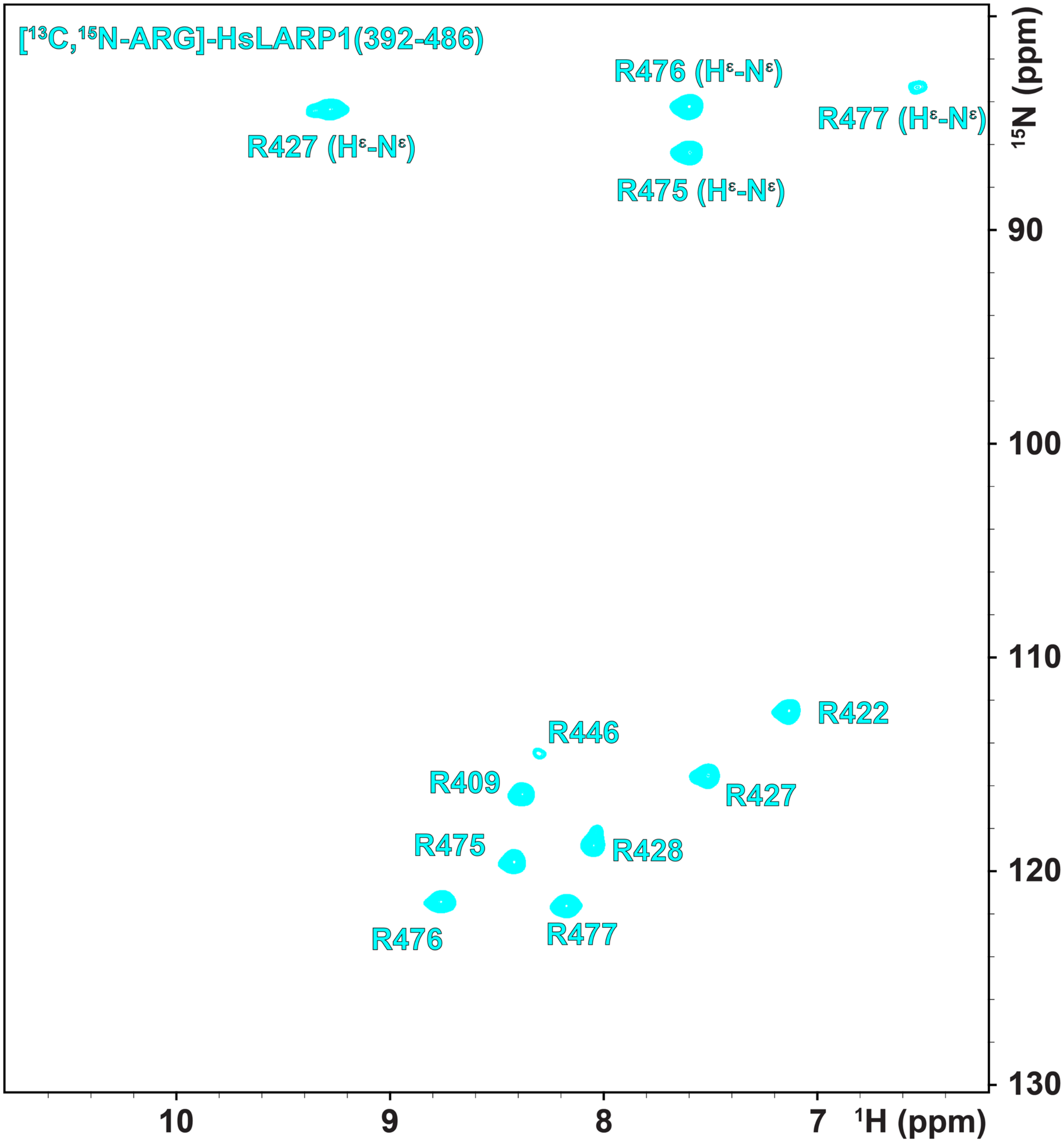


**Figure S2.** [$^1$H,$^{15}$N]-HSQC spectrum of 230 μM [$^{13}$C,$^{15}$N-Arg]-labeled HsLARP1(392-486) recorded at 700 MHz and 298 K in a buffer containing 20 mM Tris/HCl pH 7.5, 100 mM NaCl, 10% $D_2O$. Backbone and side chain resonances are indicated.

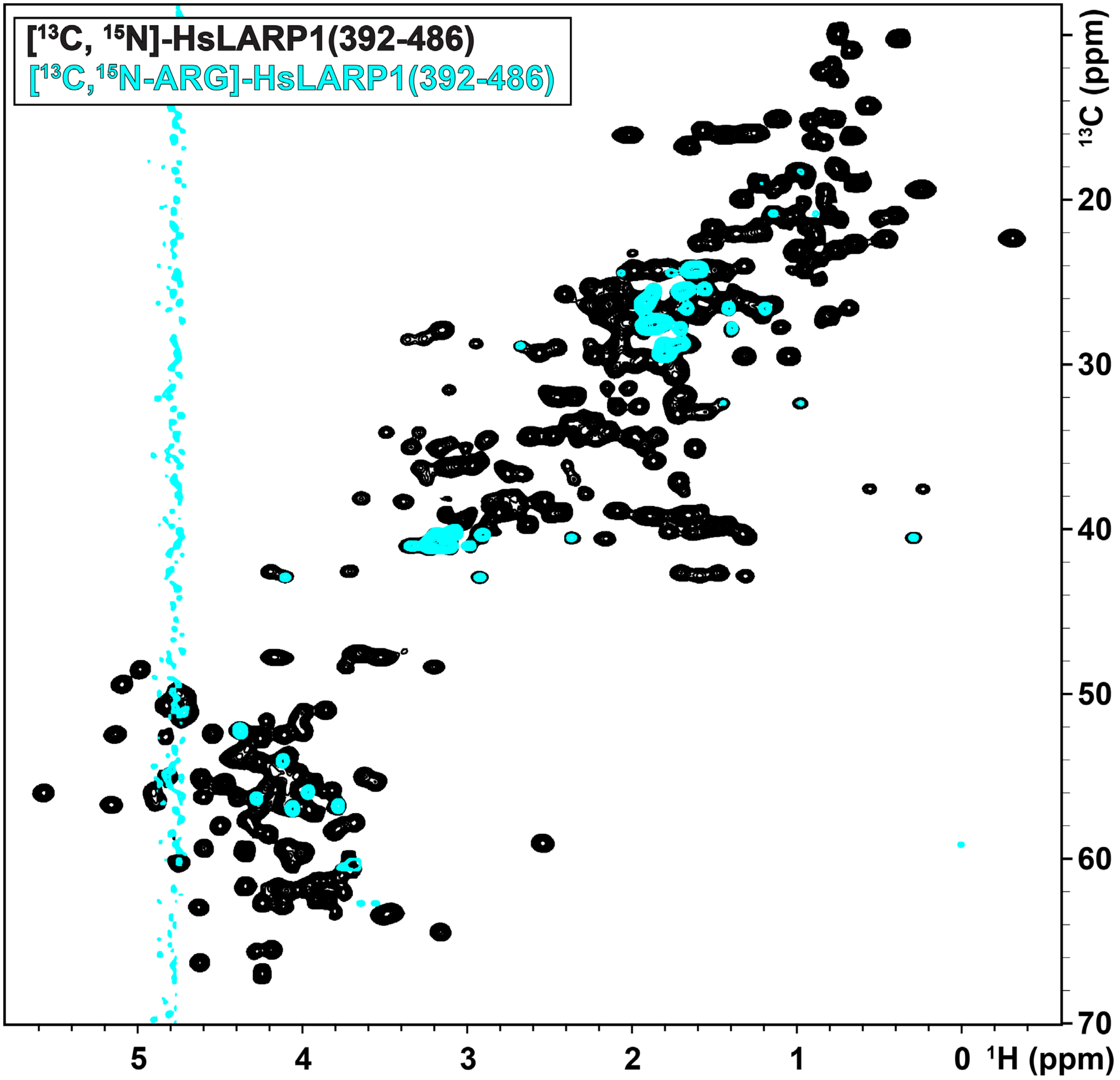


**Figure S3.** [$^1$H,$^{13}$C]-HSQC spectra of amino acid-specific labeling. The [$^1$H,$^{13}$C]-HSQC spectrum of 1.3 mM [$^{13}$C,$^{15}$N]-labeled HsLARP1(392-486) (black) and the [$^1$H,$^{13}$C]-HSQC spectrum of 230 µM [$^{13}$C,$^{15}$N-Arg]-labeled HsLARP1(392-486) (cyan) are shown. All spectra were recorded at 700 MHz and 298 K using a sample in a buffer containing 20 mM Tris/HCl pH 7.5, 100 mM NaCl, 10% $D_2O$.

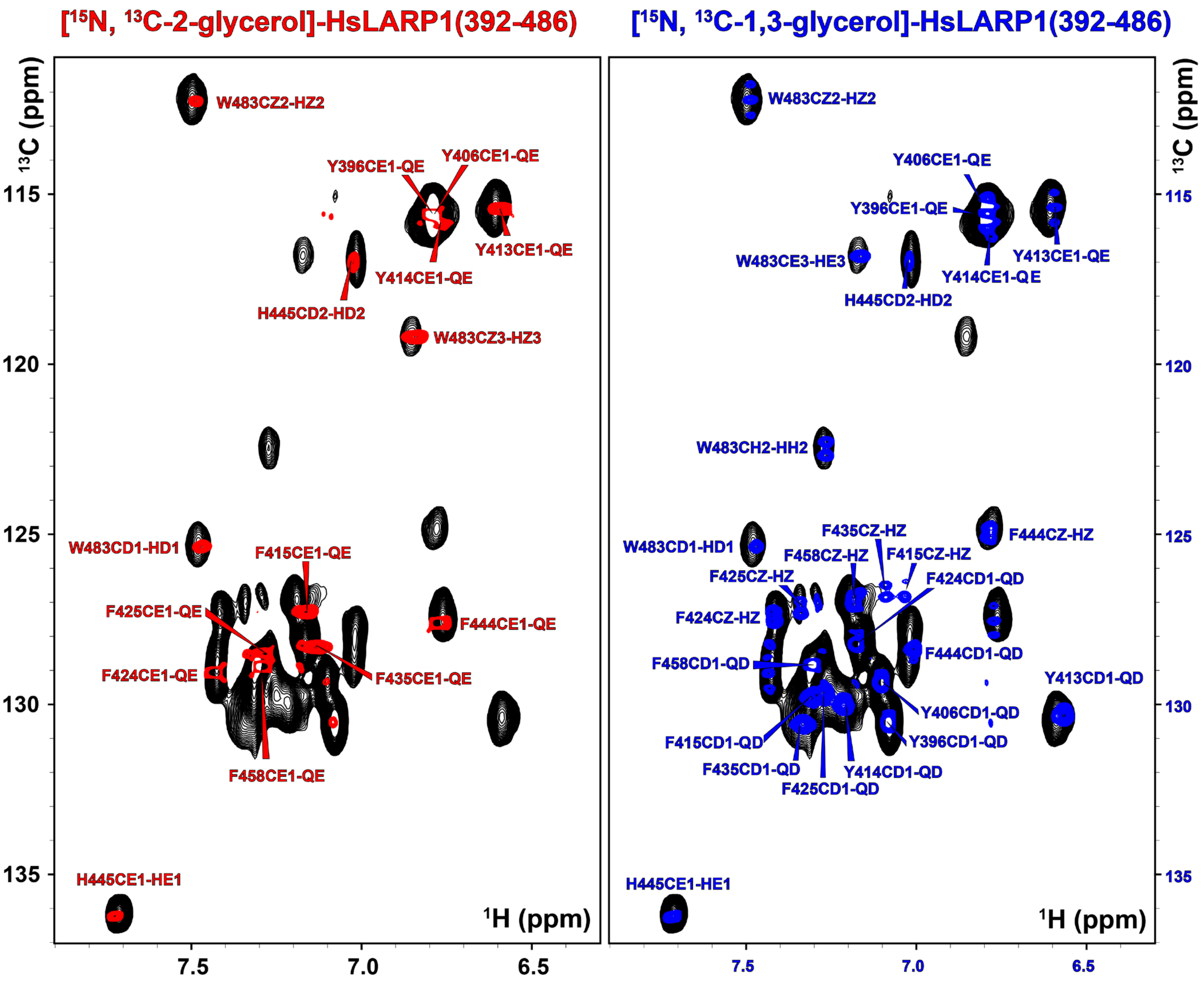


**Figure S4.** Aromatic [$^1$H,$^{13}$C]-HSQC spectra of glycerol labeling. The [$^1$H,$^{13}$C]-HSQC spectrum of 1.3 mM [$^{13}$C,$^{15}$N]-labeled HsLARP1(392-486) (black), the [$^1$H,$^{13}$C]-HSQC spectrum of 640 µM [$^{13}$C-2-glycerol,$^{15}$N]-labeled HsLARP1(392-486) (red), and the [$^1$H,$^{13}$C]-HSQC spectrum of 730 µM [$^{13}$C-1,3-glycerol,$^{15}$N]-labeled HsLARP1(392-486) (blue) are shown. All spectra were recorded at 700 MHz and 298 K using a sample in a buffer containing 20 mM Tris/HCl pH 7.5, 100 mM NaCl, 10% $D_2O$.

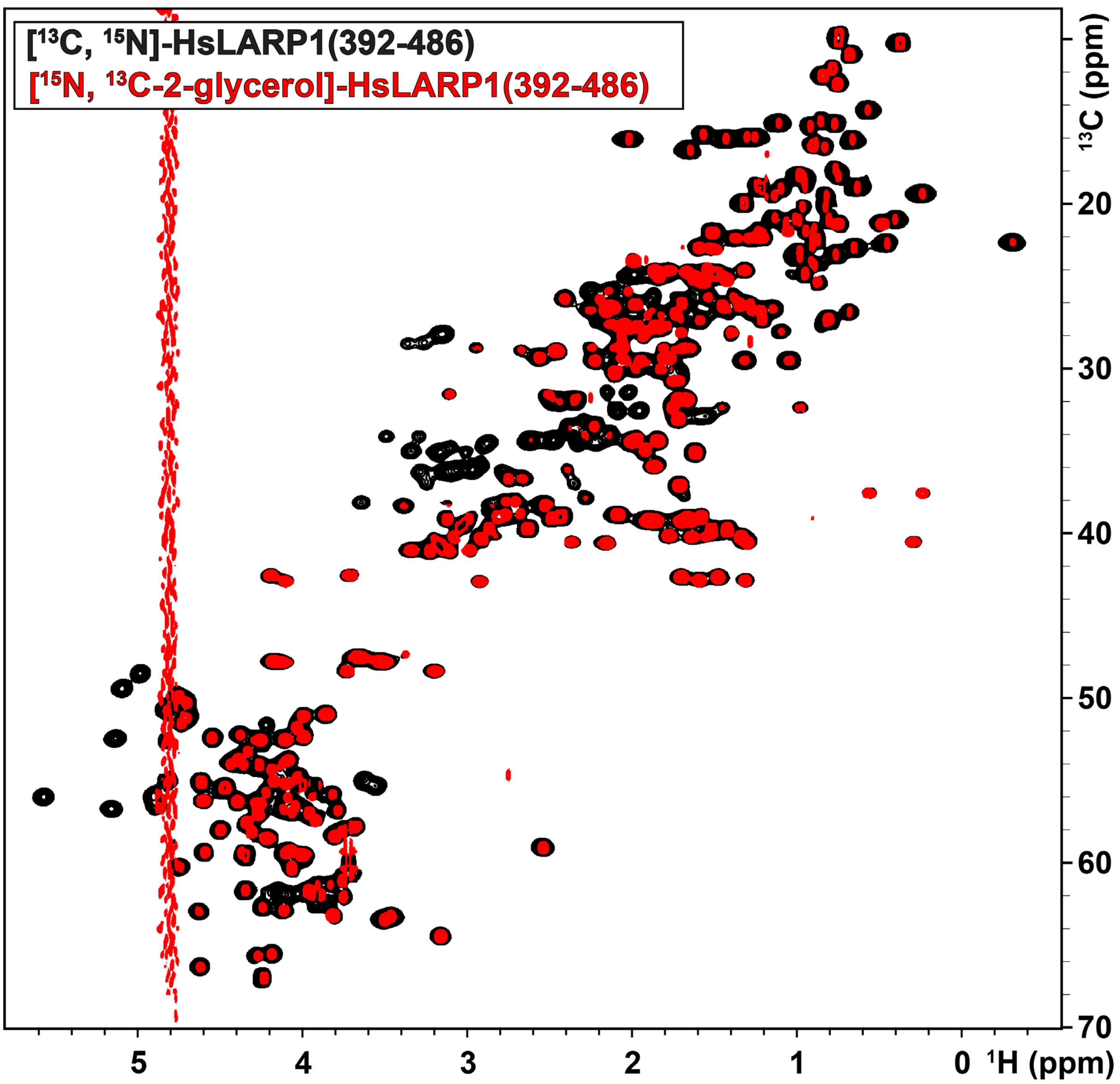


**Figure S5.** Aliphatic [$^1$H,$^{13}$C]-HSQC spectra of glycerol labeling. The [$^1$H,$^{13}$C]-HSQC spectrum of 1.3 mM [$^{13}$C,$^{15}$N]-labeled HsLARP1(392-486) (black) and the [$^1$H,$^{13}$C]-HSQC spectrum of 640 µM [$^{13}$C-2-glycerol,$^{15}$N]-labeled HsLARP1(392-486) (red) are shown. All spectra were recorded at 700 MHz and 298 K using a sample in a buffer containing 20 mM Tris/HCl pH 7.5, 100 mM NaCl, 10% $D_2O$.

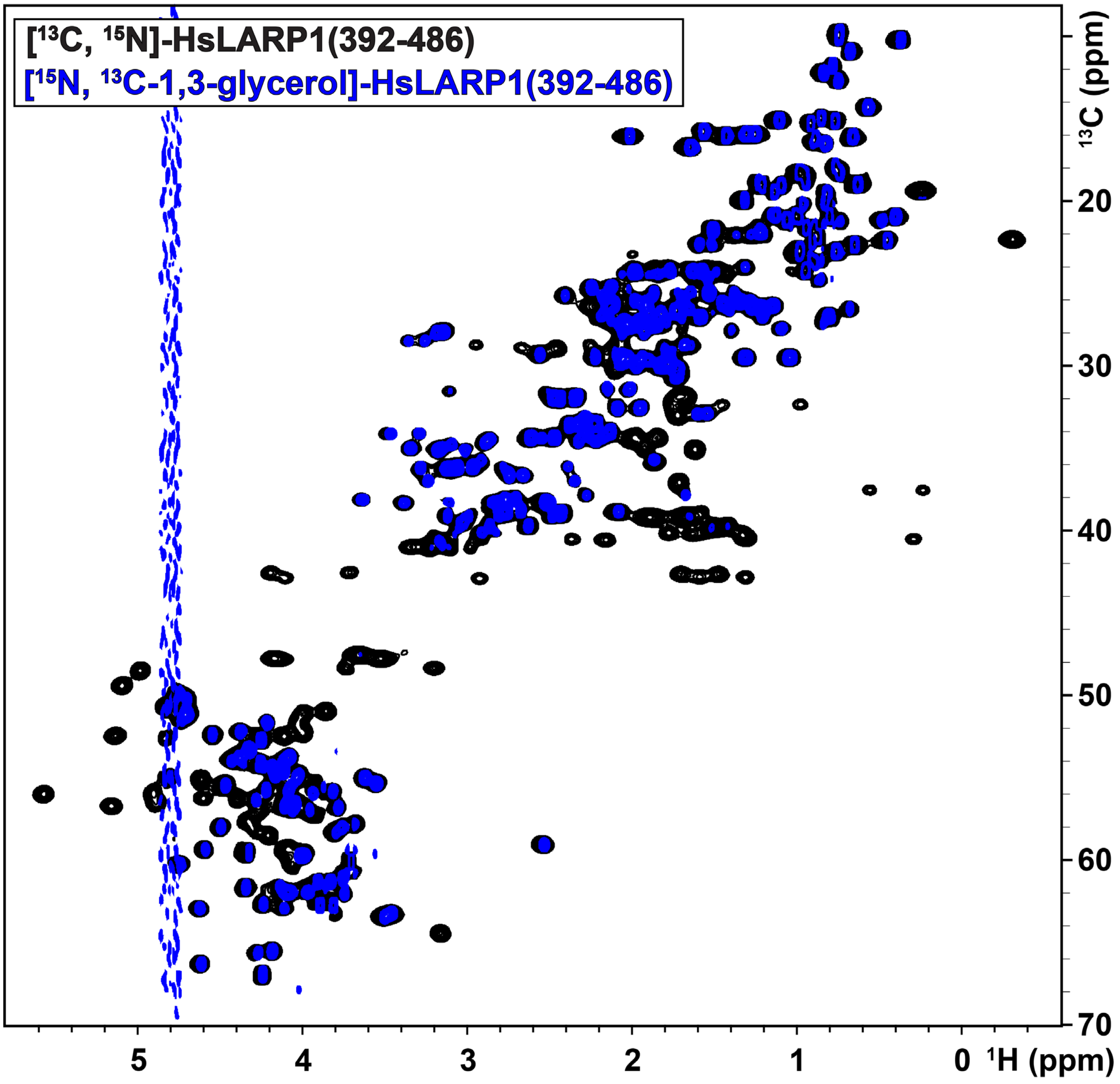


**Figure S6.** Aliphatic [$^1$H,$^{13}$C]-HSQC spectra of glycerol labeling. The [$^1$H,$^{13}$C]-HSQC spectrum of 1.3 mM [$^{13}$C,$^{15}$N]-labeled HsLARP1(392-486) (black) and the [$^1$H,$^{13}$C]-HSQC spectrum of 730 μM [$^{13}$C-1,3-glycerol,$^{15}$N]-labeled HsLARP1(392-486) (blue) are shown. All spectra were recorded at 700 MHz and 298 K using a sample in a buffer containing 20 mM Tris/HCl pH 7.5, 100 mM NaCl, 10% $D_2O$.

### Supporting Tables

**Table S1.** NMR acquisition parameters of HsLARP1(392-486). All experiments were conducted at 298 K on a 700 MHz Bruker spectrometer.

| Experiment | Labeling | ns | $t_1$ | $sw_1$ [Hz] | $o_1$ [Hz] | $t_2$ | $sw_2$ [Hz] | $o_2$ [Hz] | $t_3$ | $sw_3$ [Hz] | $o_3$ [Hz] |
|---|---|---|---|---|---|---|---|---|---|---|---|
| [$^1$H,$^{15}$N]-HSQC | [$^{13}$C,$^{15}$N] | 16 | 2048 | 10504.20 | 3291.03 | 168 | 1915.71 | 8372.42 | - | - | - |
| [$^1$H,$^{15}$N]-HSQC | [$^{15}$N-Ile] | 32 | 2048 | 10504.20 | 3292.89 | 128 | 1915.71 | 8372.42 | - | - | - |
| [$^1$H,$^{15}$N]-HSQC | [$^{15}$N-Leu] | 16 | 2048 | 10504.20 | 3290.86 | 256 | 1915.71 | 8372.42 | - | - | - |
| [$^1$H,$^{15}$N]-HSQC | [$^{15}$N-Val] | 32 | 2048 | 10504.20 | 3292.89 | 128 | 1915.71 | 8372.42 | - | - | - |
| [$^1$H,$^{15}$N]-HSQC | [$^{15}$N-Phe] | 32 | 2048 | 10504.20 | 3292.89 | 128 | 1915.71 | 8372.42 | - | - | - |
| [$^1$H,$^{15}$N]-HSQC | [$^{15}$N,$^{13}$C-Arg] | 16 | 2048 | 11160.71 | 3289.6 | 512 | 5000.00 | 6740.50 | - | - | - |
| [$^1$H, $^{13}$C]-HSQC | [$^{13}$C,$^{15}$N] | 32 | 2048 | 10504.20 | 3292.13 | 256 | 14084.51 | 7571.03 | - | - | - |
| [$^1$H, $^{13}$C]-HSQC | [$^{15}$N,$^{13}$C-Arg] | 32 | 2048 | 10504.20 | 3289.60 | 256 | 10869.57 | 6866.75 | - | - | - |
| [$^1$H, $^{13}$C]-HSQC | [$^{13}$C-1,3-glycerol,$^{15}$N] | 16 | 2048 | 10504.20 | 3292.25 | 192 | 10869.57 | 6866.75 | - | - | - |
| [$^1$H, $^{13}$C]-HSQC | [$^{13}$C-1,3-glycerol,$^{15}$N] | 32 | 2048 | 10504.20 | 3292.73 | 256 | 5000.00 | 21832.74 | - | - | - |
| [$^1$H, $^{13}$C]-HSQC | [$^{13}$C-2-glycerol,$^{15}$N] | 16 | 2048 | 10504.20 | 3292.52 | 256 | 10869.57 | 6866.75 | - | - | - |
| [$^1$H, $^{13}$C]-HSQC | [$^{13}$C-2-glycerol,$^{15}$N] | 32 | 2048 | 10504.20 | 3292.73 | 256 | 5000.00 | 21832.74 | - | - | - |
| HNCO | [$^{13}$C,$^{15}$N] | 16 | 2048 | 11160.71 | 3289.71 | 60 | 1915.71 | 30724.30 | 128 | 2958.50 | 8372.42 |
| HN(CA)CO | [$^{13}$C,$^{15}$N] | 32 | 2048 | 11160.71 | 3291.16 | 60 | 3099.38 | 30636.26 | 40 | 2128.83 | 8514.32 |
| HNCA | [$^{13}$C,$^{15}$N] | 32 | 2048 | 10504.20 | 3291.16 | 80 | 5634.50 | 9331.73 | 40 | 2127.66z | 8514.32 |
| HNCACB | [$^{13}$C,$^{15}$N] | 32 | 2048 | 11160.71 | 3289.71 | 48 | 1915.71 | 7571.07 | 128 | 14086.24 | 8372.42 |
| H(CCCO)NH | [$^{13}$C,$^{15}$N] | 16 | 2048 | 10504.20 | 3287.85 | 64 | 1915.71 | 7571.03 | 128 | 8333.33 | 8372.42 |
| (H)CC(CO)NH | [$^{13}$C,$^{15}$N] | 32 | 2048 | 11160.71 | 3291.16 | 64 | 2128.83 | 7042.82 | 192 | 10526.32 | 8514.32 |
| NOESY-[$^{15}$N]-HSQC | [$^{13}$C,$^{15}$N] | 16 | 2048 | 11235.96 | 3289.25 | 64 | 1915.71 | 8372.42 | 192 | 9803.92 | 3289.25 |
| NOESY-[$^{13}$C]-HSQC | [$^{13}$C-1,3-glycerol,$^{15}$N] | 8 | 2048 | 9803.92 | 3292.25 | 128 | 10869.57 | 6866.75 | 192 | 9803.92 | 3292.25 |
| NOESY-[$^{13}$C]-HSQC | [$^{13}$C-1,3-glycerol,$^{15}$N] | 16 | 2048 | 9803.92 | 3292.73 | 64 | 5000.00 | 21832.74 | 192 | 9803.92 | 3292.73 |
| NOESY-[$^{13}$C]-HSQC | [$^{13}$C-2-glycerol,$^{15}$N] | 8 | 2048 | 9803.92 | 3292.52 | 128 | 10869.57 | 6866.75 | 192 | 9803.92 | 3292.52 |
| NOESY-[$^{13}$C]-HSQC | [$^{13}$C-2-glycerol,$^{15}$N] | 16 | 2048 | 9803.92 | 3288.65 | 64 | 5000.00 | 21832.74 | 192 | 9803.92 | 3288.65 |

**Table S2.** Chemical shift assignment of HsLARP1(392-486)

| Residue | Atom | Chemical Shift [ppm] | Standard Deviation [ppm] |
|---|---|---|---|
| S392 | CA | 55.292 | 0.010 |
| S392 | CB | 62.750 | 0.006 |
| S392 | HA | 3.998 | 0.001 |
| S392 | HB2 | 3.892 | 0.001 |
| S392 | HB3 | 3.812 | 0.000 |
| T393 | C | 172.000 | 0.001 |
| T393 | CA | 59.563 | 0.021 |
| T393 | CB | 66.979 | 0.022 |
| T393 | CG2 | 19.001 | 0.040 |
| T393 | HA | 4.335 | 0.010 |
| T393 | HB | 4.241 | 0.004 |
| T393 | QG2 | 1.210 | 0.003 |
| E394 | C | 173.686 | 0.044 |
| E394 | CA | 54.135 | 0.053 |
| E394 | CB | 27.442 | 0.028 |
| E394 | CG | 33.530 | 0.009 |
| E394 | H | 8.547 | 0.006 |
| E394 | HA | 4.274 | 0.009 |
| E394 | HB2 | 1.962 | 0.002 |
| E394 | HB3 | 1.864 | 0.004 |
| E394 | N | 123.503 | 0.014 |
| E394 | QG | 2.220 | 0.008 |
| L395 | C | 174.336 | 0.012 |
| L395 | CA | 52.793 | 0.051 |
| L395 | CB | 39.807 | 0.029 |
| L395 | CD1 | 22.169 | 0.001 |
| L395 | CD2 | 20.959 | 0.029 |
| L395 | CG | 24.257 | 0.000 |
| L395 | H | 8.167 | 0.017 |
| L395 | HA | 4.257 | 0.009 |
| L395 | HB2 | 1.521 | 0.006 |
| L395 | HB3 | 1.424 | 0.005 |
| L395 | HG | 1.484 | 0.013 |
| L395 | N | 122.852 | 0.085 |
| L395 | QD1 | 0.875 | 0.002 |
| L395 | QD2 | 0.810 | 0.006 |
| Y396 | C | 173.102 | 0.033 |
| Y396 | CA | 55.167 | 0.044 |
| Y396 | CB | 36.149 | 0.047 |
| Y396 | CD1 | 130.513 | 0.013 |
| Y396 | CE1 | 115.593 | 0.020 |
| Y396 | H | 8.076 | 0.011 |
| Y396 | HA | 4.615 | 0.004 |
| Y396 | HB2 | 3.060 | 0.001 |
| Y396 | HB3 | 2.968 | 0.002 |
| Y396 | N | 120.083 | 0.009 |
| Y396 | QD | 7.084 | 0.002 |
| Y396 | QE | 6.793 | 0.004 |
| S397 | C | 171.883 | 0.017 |
| S397 | CA | 55.526 | 0.052 |
| S397 | CB | 61.361 | 0.071 |
| S397 | H | 8.169 | 0.007 |
| S397 | HA | 4.477 | 0.013 |
| S397 | HB2 | 3.910 | 0.002 |
| S397 | HB3 | 3.839 | 0.017 |
| S397 | N | 117.239 | 0.032 |

| Residue | Atom | Chemical Shift [ppm] | Standard Deviation [ppm] |
|---|---|---|---|
| V398 | C | 173.391 | 0.018 |
| V398 | CA | 60.308 | 0.028 |
| V398 | CB | 30.211 | 0.048 |
| V398 | CG1 | 18.285 | 0.046 |
| V398 | H | 8.099 | 0.011 |
| V398 | HA | 4.077 | 0.009 |
| V398 | HB | 2.108 | 0.007 |
| V398 | N | 121.532 | 0.046 |
| V398 | QQG | 0.978 | 0.008 |
| D399 | C | 174.694 | 0.009 |
| D399 | CA | 52.426 | 0.042 |
| D399 | CB | 38.807 | 0.055 |
| D399 | H | 8.240 | 0.006 |
| D399 | HA | 4.554 | 0.010 |
| D399 | HB2 | 2.777 | 0.009 |
| D399 | HB3 | 2.681 | 0.011 |
| D399 | N | 122.718 | 0.029 |
| Q400 | C | 175.089 | 0.006 |
| Q400 | CA | 56.030 | 0.035 |
| Q400 | CB | 26.508 | 0.059 |
| Q400 | CD | 177.214 | 0.006 |
| Q400 | CG | 32.016 | 0.038 |
| Q400 | H | 8.557 | 0.010 |
| Q400 | HA | 4.092 | 0.012 |
| Q400 | HE21 | 7.551 | 0.004 |
| Q400 | HE22 | 6.843 | 0.003 |
| Q400 | HG2 | 2.429 | 0.013 |
| Q400 | HG3 | 2.356 | 0.007 |
| Q400 | N | 122.357 | 0.033 |
| Q400 | NE2 | 112.191 | 0.024 |
| Q400 | QB | 2.140 | 0.009 |
| E401 | C | 176.605 | 0.000 |
| E401 | CA | 56.667 | 0.015 |
| E401 | CB | 26.322 | 0.040 |
| E401 | CG | 33.591 | 0.029 |
| E401 | H | 8.323 | 0.008 |
| E401 | HA | 4.068 | 0.018 |
| E401 | HG2 | 2.367 | 0.012 |
| E401 | HG3 | 2.338 | 0.003 |
| E401 | N | 119.035 | 0.011 |
| E401 | QB | 2.116 | 0.005 |
| L402 | C | 175.921 | 0.002 |
| L402 | CA | 54.648 | 0.086 |
| L402 | CB | 39.055 | 0.075 |
| L402 | CD1 | 21.658 | 0.044 |
| L402 | CG | 24.187 | 0.069 |
| L402 | H | 7.843 | 0.007 |
| L402 | HA | 4.108 | 0.004 |
| L402 | HB2 | 1.663 | 0.000 |
| L402 | HB3 | 1.579 | 0.001 |
| L402 | HG | 1.499 | 0.004 |
| L402 | N | 121.713 | 0.019 |
| L402 | QQD | 0.898 | 0.006 |
| L403 | C | 176.287 | 0.018 |
| L403 | CA | 55.607 | 0.092 |
| L403 | CB | 39.222 | 0.029 |
| L403 | CD1 | 22.981 | 0.084 |
| L403 | CD2 | 21.135 | 0.012 |

| Residue | Atom | Chemical Shift [ppm] | Standard Deviation [ppm] |
|---|---|---|---|
| L403 | CG | 24.523 | 0.027 |
| L403 | H | 7.878 | 0.008 |
| L403 | HA | 4.086 | 0.006 |
| L403 | HB2 | 1.862 | 0.010 |
| L403 | HB3 | 1.666 | 0.003 |
| L403 | HG | 1.629 | 0.007 |
| L403 | N | 119.598 | 0.019 |
| L403 | QD1 | 0.981 | 0.004 |
| L403 | QD2 | 1.060 | 0.002 |
| K404 | C | 175.282 | 0.011 |
| K404 | CA | 58.440 | 0.047 |
| K404 | CB | 29.660 | 0.026 |
| K404 | CD | 27.041 | 0.032 |
| K404 | CE | 39.010 | 0.000 |
| K404 | CG | 24.032 | 0.040 |
| K404 | H | 8.184 | 0.007 |
| K404 | HA | 3.804 | 0.010 |
| K404 | HB2 | 2.013 | 0.004 |
| K404 | HB3 | 1.903 | 0.001 |
| K404 | HD2 | 1.709 | 0.002 |
| K404 | HD3 | 1.586 | 0.002 |
| K404 | HG2 | 1.771 | 0.003 |
| K404 | HG3 | 1.318 | 0.004 |
| K404 | N | 116.733 | 0.011 |
| K404 | QE | 2.806 | 0.010 |
| D405 | C | 175.780 | 0.009 |
| D405 | CA | 55.517 | 0.066 |
| D405 | CB | 38.962 | 0.030 |
| D405 | H | 7.663 | 0.004 |
| D405 | HA | 4.470 | 0.004 |
| D405 | N | 119.333 | 0.024 |
| D405 | QB | 2.791 | 0.009 |
| Y406 | C | 176.821 | 0.002 |
| Y406 | CA | 57.615 | 0.090 |
| Y406 | CB | 35.014 | 0.008 |
| Y406 | CD1 | 129.340 | 0.018 |
| Y406 | CE1 | 115.593 | 0.020 |
| Y406 | H | 8.309 | 0.006 |
| Y406 | HA | 4.326 | 0.010 |
| Y406 | HB2 | 3.344 | 0.013 |
| Y406 | HB3 | 3.167 | 0.012 |
| Y406 | N | 119.330 | 0.023 |
| Y406 | QD | 7.103 | 0.004 |
| Y406 | QE | 6.800 | 0.006 |
| I407 | C | 174.386 | 0.005 |
| I407 | CA | 63.266 | 0.063 |
| I407 | CB | 35.880 | 0.032 |
| I407 | CD1 | 12.724 | 0.031 |
| I407 | CG1 | 27.448 | 0.053 |
| I407 | CG2 | 16.521 | 0.000 |
| I407 | H | 8.803 | 0.004 |
| I407 | HA | 3.466 | 0.006 |
| I407 | HB | 1.873 | 0.004 |
| I407 | HG12 | 2.016 | 0.001 |
| I407 | HG13 | 0.838 | 0.005 |
| I407 | N | 120.645 | 0.026 |
| I407 | QD1 | 0.753 | 0.003 |
| I407 | QG2 | 0.828 | 0.005 |

| Residue | Atom | Chemical Shift [ppm] | Standard Deviation [ppm] |
|---|---|---|---|
| K408 | C | 175.317 | 0.013 |
| K408 | CA | 57.854 | 0.036 |
| K408 | CB | 29.345 | 0.037 |
| K408 | CD | 26.820 | 0.012 |
| K408 | CE | 39.139 | 0.015 |
| K408 | CG | 22.051 | 0.039 |
| K408 | H | 8.534 | 0.004 |
| K408 | HA | 3.693 | 0.009 |
| K408 | HD2 | 1.898 | 0.001 |
| K408 | HD3 | 1.823 | 0.001 |
| K408 | HE2 | 3.121 | 0.006 |
| K408 | HE3 | 2.993 | 0.002 |
| K408 | HG2 | 1.368 | 0.005 |
| K408 | HG3 | 1.214 | 0.006 |
| K408 | N | 119.815 | 0.035 |
| K408 | QB | 2.035 | 0.008 |
| R409 | C | 177.000 | 0.031 |
| R409 | CA | 57.003 | 0.033 |
| R409 | CB | 27.701 | 0.094 |
| R409 | CD | 41.039 | 0.014 |
| R409 | CG | 25.593 | 0.058 |
| R409 | H | 8.383 | 0.007 |
| R409 | HA | 4.067 | 0.011 |
| R409 | HD2 | 3.207 | 0.014 |
| R409 | HD3 | 3.118 | 0.006 |
| R409 | HG2 | 1.869 | 0.002 |
| R409 | HG3 | 1.710 | 0.004 |
| R409 | N | 116.397 | 0.019 |
| R409 | QB | 1.886 | 0.003 |
| Q410 | C | 174.880 | 0.026 |
| Q410 | CA | 55.122 | 0.053 |
| Q410 | CB | 26.052 | 0.054 |
| Q410 | CD | 175.946 | 0.000 |
| Q410 | CG | 31.351 | 0.013 |
| Q410 | H | 8.118 | 0.010 |
| Q410 | HA | 4.053 | 0.010 |
| Q410 | HB2 | 2.159 | 0.003 |
| Q410 | HB3 | 1.983 | 0.005 |
| Q410 | HE21 | 6.969 | 0.011 |
| Q410 | HE22 | 6.400 | 0.006 |
| Q410 | HG2 | 2.151 | 0.009 |
| Q410 | HG3 | 2.010 | 0.005 |
| Q410 | N | 118.616 | 0.014 |
| Q410 | NE2 | 114.176 | 0.019 |
| I411 | C | 175.687 | 0.023 |
| I411 | CA | 62.927 | 0.067 |
| I411 | CB | 34.560 | 0.053 |
| I411 | CD1 | 12.221 | 0.002 |
| I411 | CG1 | 27.724 | 0.072 |
| I411 | CG2 | 15.093 | 0.041 |
| I411 | H | 8.312 | 0.005 |
| I411 | HA | 4.121 | 0.010 |
| I411 | HB | 1.993 | 0.006 |
| I411 | HG12 | 2.015 | 0.001 |
| I411 | HG13 | 1.099 | 0.007 |
| I411 | N | 117.821 | 0.028 |
| I411 | QD1 | 0.842 | 0.013 |
| I411 | QG2 | 1.108 | 0.004 |

| Residue | Atom | Chemical Shift [ppm] | Standard Deviation [ppm] |
|---|---|---|---|
| E412 | C | 176.742 | 0.043 |
| E412 | CA | 58.073 | 0.060 |
| E412 | CB | 26.405 | 0.063 |
| E412 | CG | 35.702 | 0.046 |
| E412 | H | 9.057 | 0.005 |
| E412 | HA | 3.764 | 0.006 |
| E412 | HB2 | 2.252 | 0.006 |
| E412 | HB3 | 1.878 | 0.007 |
| E412 | HG2 | 2.915 | 0.001 |
| E412 | HG3 | 1.867 | 0.001 |
| E412 | N | 117.854 | 0.024 |
| Y413 | C | 176.173 | 0.004 |
| Y413 | CA | 59.382 | 0.061 |
| Y413 | CB | 34.689 | 0.095 |
| Y413 | CD1 | 130.313 | 0.012 |
| Y413 | CE1 | 115.398 | 0.021 |
| Y413 | H | 7.656 | 0.006 |
| Y413 | HA | 4.097 | 0.008 |
| Y413 | HB2 | 3.096 | 0.009 |
| Y413 | HB3 | 2.899 | 0.010 |
| Y413 | N | 118.711 | 0.021 |
| Y413 | QD | 6.571 | 0.004 |
| Y413 | QE | 6.594 | 0.003 |
| Y414 | C | 173.130 | 0.022 |
| Y414 | CA | 57.517 | 0.087 |
| Y414 | CB | 34.386 | 0.051 |
| Y414 | CD1 | 130.015 | 0.020 |
| Y414 | CE1 | 115.807 | 0.027 |
| Y414 | H | 7.427 | 0.005 |
| Y414 | HA | 3.932 | 0.007 |
| Y414 | HB2 | 2.871 | 0.008 |
| Y414 | HB3 | 2.544 | 0.007 |
| Y414 | N | 117.859 | 0.016 |
| Y414 | QD | 7.217 | 0.005 |
| Y414 | QE | 6.785 | 0.010 |
| F415 | C | 170.598 | 0.012 |
| F415 | CA | 55.229 | 0.077 |
| F415 | CB | 37.015 | 0.078 |
| F415 | CD1 | 129.813 | 0.018 |
| F415 | CE1 | 127.279 | 0.018 |
| F415 | CZ | 126.620 | 0.000 |
| F415 | H | 7.095 | 0.005 |
| F415 | HA | 4.112 | 0.012 |
| F415 | HB2 | 3.239 | 0.012 |
| F415 | HB3 | 2.349 | 0.008 |
| F415 | HZ | 7.035 | 0.006 |
| F415 | N | 110.642 | 0.033 |
| F415 | QD | 7.308 | 0.005 |
| F415 | QE | 7.164 | 0.006 |
| S416 | C | 171.581 | 0.019 |
| S416 | CA | 55.077 | 0.077 |
| S416 | CB | 61.966 | 0.064 |
| S416 | H | 7.391 | 0.010 |
| S416 | HA | 4.162 | 0.018 |
| S416 | N | 116.504 | 0.013 |
| S416 | QB | 4.114 | 0.012 |
| V417 | C | 174.192 | 0.021 |
| V417 | CA | 64.463 | 0.080 |

| Residue | Atom | Chemical Shift [ppm] | Standard Deviation [ppm] |
|---|---|---|---|
| V417 | CB | 28.627 | 0.051 |
| V417 | CG1 | 19.419 | 0.084 |
| V417 | CG2 | 19.085 | 0.029 |
| V417 | H | 8.808 | 0.008 |
| V417 | HA | 3.161 | 0.002 |
| V417 | HB | 2.056 | 0.005 |
| V417 | N | 123.023 | 0.033 |
| V417 | QG1 | 1.132 | 0.005 |
| V417 | QG2 | 1.099 | 0.013 |
| V417 | QQG | 1.110 | 0.007 |
| D418 | C | 175.381 | 0.004 |
| D418 | CA | 54.348 | 0.069 |
| D418 | CB | 38.292 | 0.044 |
| D418 | H | 7.936 | 0.005 |
| D418 | HA | 4.178 | 0.010 |
| D418 | N | 115.216 | 0.010 |
| D418 | QB | 2.526 | 0.004 |
| N419 | C | 174.780 | 0.001 |
| N419 | CA | 54.059 | 0.055 |
| N419 | CB | 37.837 | 0.055 |
| N419 | H | 7.299 | 0.006 |
| N419 | HA | 4.365 | 0.007 |
| N419 | HB2 | 2.281 | 0.004 |
| N419 | HB3 | 1.678 | 0.008 |
| N419 | HD21 | 8.719 | 0.006 |
| N419 | HD22 | 8.271 | 0.005 |
| N419 | N | 113.599 | 0.033 |
| N419 | ND2 | 120.215 | 0.014 |
| L420 | C | 175.298 | 0.005 |
| L420 | CA | 55.356 | 0.077 |
| L420 | CB | 37.567 | 0.056 |
| L420 | CD1 | 22.346 | 0.013 |
| L420 | CD2 | 20.951 | 0.014 |
| L420 | CG | 22.861 | 0.002 |
| L420 | H | 8.478 | 0.003 |
| L420 | HA | 3.554 | 0.006 |
| L420 | HB2 | 0.554 | 0.009 |
| L420 | HB3 | 0.241 | 0.008 |
| L420 | HG | 1.503 | 0.005 |
| L420 | N | 119.194 | 0.039 |
| L420 | QD1 | -0.314 | 0.005 |
| L420 | QD2 | 0.408 | 0.009 |
| E421 | C | 174.314 | 0.007 |
| E421 | CA | 56.765 | 0.041 |
| E421 | CB | 27.524 | 0.048 |
| E421 | CG | 34.362 | 0.019 |
| E421 | H | 8.102 | 0.005 |
| E421 | HA | 4.113 | 0.006 |
| E421 | HG2 | 2.617 | 0.006 |
| E421 | HG3 | 2.473 | 0.003 |
| E421 | N | 114.840 | 0.020 |
| E421 | QB | 2.049 | 0.009 |
| R422 | C | 172.840 | 0.072 |
| R422 | CA | 52.278 | 0.072 |
| R422 | CB | 29.372 | 0.064 |
| R422 | CD | 40.595 | 0.028 |
| R422 | CG | 24.248 | 0.027 |
| R422 | H | 7.136 | 0.006 |

| Residue | Atom | Chemical Shift [ppm] | Standard Deviation [ppm] |
|---|---|---|---|
| R422 | HA | 4.376 | 0.006 |
| R422 | HG2 | 1.627 | 0.004 |
| R422 | HG3 | 1.585 | 0.010 |
| R422 | N | 112.424 | 0.021 |
| R422 | QB | 1.785 | 0.011 |
| R422 | QD | 3.166 | 0.006 |
| D423 | C | 172.808 | 0.019 |
| D423 | CA | 49.956 | 0.065 |
| D423 | CB | 36.164 | 0.082 |
| D423 | H | 7.720 | 0.006 |
| D423 | HA | 4.742 | 0.008 |
| D423 | HB2 | 2.780 | 0.003 |
| D423 | HB3 | 2.394 | 0.011 |
| D423 | N | 122.797 | 0.015 |
| F424 | C | 174.686 | 0.026 |
| F424 | CA | 57.148 | 0.034 |
| F424 | CB | 35.203 | 0.043 |
| F424 | CD1 | 128.211 | 0.016 |
| F424 | CE1 | 129.057 | 0.010 |
| F424 | CZ | 127.437 | 0.010 |
| F424 | H | 7.727 | 0.009 |
| F424 | HA | 4.275 | 0.007 |
| F424 | HB2 | 3.174 | 0.007 |
| F424 | HB3 | 3.010 | 0.014 |
| F424 | HZ | 7.420 | 0.002 |
| F424 | N | 122.999 | 0.022 |
| F424 | QD | 7.179 | 0.003 |
| F424 | QE | 7.433 | 0.003 |
| F425 | C | 174.654 | 0.019 |
| F425 | CA | 58.564 | 0.080 |
| F425 | CB | 36.211 | 0.028 |
| F425 | CD1 | 129.659 | 0.008 |
| F425 | CE1 | 128.501 | 0.014 |
| F425 | CZ | 127.312 | 0.005 |
| F425 | H | 8.316 | 0.005 |
| F425 | HA | 4.220 | 0.007 |
| F425 | HB2 | 3.287 | 0.010 |
| F425 | HB3 | 3.130 | 0.011 |
| F425 | HZ | 7.327 | 0.011 |
| F425 | N | 119.031 | 0.007 |
| F425 | QD | 7.274 | 0.003 |
| F425 | QE | 7.288 | 0.004 |
| L426 | C | 176.867 | 0.039 |
| L426 | CA | 55.058 | 0.085 |
| L426 | CB | 40.180 | 0.038 |
| L426 | CD1 | 22.621 | 0.075 |
| L426 | CD2 | 22.405 | 0.009 |
| L426 | CG | 23.970 | 0.005 |
| L426 | H | 7.830 | 0.015 |
| L426 | HA | 3.621 | 0.015 |
| L426 | HB2 | 1.630 | 0.008 |
| L426 | HB3 | 1.335 | 0.004 |
| L426 | HG | 1.542 | 0.003 |
| L426 | N | 119.441 | 0.035 |
| L426 | QD1 | 0.654 | 0.005 |
| L426 | QD2 | 0.451 | 0.006 |
| R427 | C | 177.197 | 0.017 |
| R427 | CA | 56.367 | 0.070 |

| Residue | Atom | Chemical Shift [ppm] | Standard Deviation [ppm] |
|---|---|---|---|
| R427 | CB | 28.847 | 0.048 |
| R427 | CD | 42.931 | 0.040 |
| R427 | CG | 24.437 | 0.033 |
| R427 | CZ | 157.185 | 0.000 |
| R427 | H | 7.518 | 0.012 |
| R427 | HA | 4.280 | 0.004 |
| R427 | HB2 | 2.674 | 0.005 |
| R427 | HB3 | 1.738 | 0.011 |
| R427 | HD2 | 4.109 | 0.005 |
| R427 | HD3 | 2.925 | 0.003 |
| R427 | HE | 9.281 | 0.009 |
| R427 | HG2 | 2.060 | 0.002 |
| R427 | HG3 | 1.761 | 0.006 |
| R427 | N | 115.449 | 0.023 |
| R427 | NE | 84.357 | 0.014 |
| R428 | C | 175.262 | 0.007 |
| R428 | CA | 55.119 | 0.076 |
| R428 | CB | 27.701 | 0.025 |
| R428 | CD | 40.955 | 0.007 |
| R428 | CG | 26.539 | 0.034 |
| R428 | H | 8.049 | 0.004 |
| R428 | HA | 4.806 | 0.012 |
| R428 | HG2 | 1.935 | 0.003 |
| R428 | HG3 | 1.656 | 0.002 |
| R428 | N | 118.806 | 0.026 |
| R428 | QB | 1.901 | 0.012 |
| R428 | QD | 3.154 | 0.003 |
| K429 | C | 173.886 | 0.014 |
| K429 | CA | 52.584 | 0.058 |
| K429 | CB | 30.123 | 0.014 |
| K429 | CD | 25.657 | 0.043 |
| K429 | CE | 39.740 | 0.031 |
| K429 | CG | 21.693 | 0.051 |
| K429 | H | 7.258 | 0.007 |
| K429 | HA | 4.256 | 0.005 |
| K429 | HB2 | 1.800 | 0.003 |
| K429 | HB3 | 1.729 | 0.005 |
| K429 | HD2 | 1.537 | 0.004 |
| K429 | HD3 | 1.375 | 0.010 |
| K429 | HE2 | 2.866 | 0.005 |
| K429 | HE3 | 2.637 | 0.012 |
| K429 | N | 116.638 | 0.022 |
| K429 | QG | 1.223 | 0.003 |
| M430 | C | 174.194 | 0.017 |
| M430 | CA | 54.070 | 0.070 |
| M430 | CB | 31.607 | 0.053 |
| M430 | CE | 16.056 | 0.008 |
| M430 | CG | 31.550 | 0.012 |
| M430 | H | 7.000 | 0.004 |
| M430 | HA | 4.431 | 0.003 |
| M430 | HG2 | 3.102 | 0.007 |
| M430 | HG3 | 2.502 | 0.006 |
| M430 | N | 119.053 | 0.023 |
| M430 | QB | 2.328 | 0.005 |
| M430 | QE | 2.017 | 0.003 |
| D431 | C | 174.803 | 0.011 |
| D431 | CA | 49.968 | 0.039 |
| D431 | CB | 38.305 | 0.029 |

| Residue | Atom | Chemical Shift [ppm] | Standard Deviation [ppm] |
|---|---|---|---|
| D431 | H | 8.330 | 0.005 |
| D431 | HA | 4.755 | 0.002 |
| D431 | HB2 | 3.386 | 0.002 |
| D431 | HB3 | 2.846 | 0.006 |
| D431 | N | 120.598 | 0.024 |
| A432 | C | 176.252 | 0.023 |
| A432 | CA | 52.395 | 0.033 |
| A432 | CB | 16.047 | 0.031 |
| A432 | H | 8.276 | 0.005 |
| A432 | HA | 4.000 | 0.010 |
| A432 | N | 117.726 | 0.013 |
| A432 | QB | 1.429 | 0.001 |
| D433 | C | 172.610 | 0.030 |
| D433 | CA | 51.581 | 0.087 |
| D433 | CB | 40.100 | 0.028 |
| D433 | H | 8.040 | 0.004 |
| D433 | HA | 4.742 | 0.007 |
| D433 | HB2 | 2.900 | 0.010 |
| D433 | HB3 | 2.825 | 0.003 |
| D433 | N | 115.245 | 0.021 |
| G434 | C | 171.129 | 0.018 |
| G434 | CA | 42.574 | 0.032 |
| G434 | H | 8.373 | 0.008 |
| G434 | HA2 | 4.196 | 0.002 |
| G434 | HA3 | 3.713 | 0.004 |
| G434 | N | 106.196 | 0.014 |
| F435 | C | 174.823 | 0.019 |
| F435 | CA | 56.664 | 0.065 |
| F435 | CB | 38.111 | 0.043 |
| F435 | CD1 | 130.587 | 0.017 |
| F435 | CE1 | 128.309 | 0.020 |
| F435 | CZ | 126.650 | 0.016 |
| F435 | H | 8.403 | 0.004 |
| F435 | HA | 4.890 | 0.010 |
| F435 | HB2 | 3.646 | 0.005 |
| F435 | HB3 | 2.820 | 0.014 |
| F435 | HZ | 7.092 | 0.008 |
| F435 | N | 117.589 | 0.016 |
| F435 | QD | 7.332 | 0.006 |
| F435 | QE | 7.135 | 0.003 |
| L436 | C | 171.434 | 0.000 |
| L436 | CA | 49.497 | 0.047 |
| L436 | CB | 42.662 | 0.011 |
| L436 | CD1 | 24.229 | 0.015 |
| L436 | CD2 | 22.931 | 0.078 |
| L436 | CG | 24.379 | 0.048 |
| L436 | H | 9.241 | 0.004 |
| L436 | HA | 5.093 | 0.010 |
| L436 | HB2 | 1.698 | 0.004 |
| L436 | HB3 | 1.476 | 0.012 |
| L436 | HG | 1.841 | 0.004 |
| L436 | N | 118.600 | 0.016 |
| L436 | QD1 | 0.958 | 0.008 |
| L436 | QD2 | 0.985 | 0.008 |
| P437 | C | 176.406 | 0.013 |
| P437 | CA | 60.220 | 0.096 |
| P437 | CB | 29.335 | 0.032 |
| P437 | CD | 47.773 | 0.007 |

| Residue | Atom | Chemical Shift [ppm] | Standard Deviation [ppm] |
|---|---|---|---|
| P437 | CG | 25.241 | 0.031 |
| P437 | HA | 4.752 | 0.006 |
| P437 | HB2 | 2.560 | 0.004 |
| P437 | HB3 | 2.078 | 0.012 |
| P437 | HD2 | 4.177 | 0.003 |
| P437 | HD3 | 3.530 | 0.006 |
| P437 | HG2 | 2.246 | 0.003 |
| P437 | HG3 | 2.125 | 0.002 |
| I438 | C | 173.791 | 0.015 |
| I438 | CA | 61.100 | 0.086 |
| I438 | CB | 35.124 | 0.031 |
| I438 | CD1 | 10.231 | 0.036 |
| I438 | CG1 | 27.008 | 0.019 |
| I438 | CG2 | 15.121 | 0.020 |
| I438 | H | 8.826 | 0.005 |
| I438 | HA | 3.761 | 0.006 |
| I438 | HB | 1.617 | 0.004 |
| I438 | HG12 | 1.215 | 0.011 |
| I438 | HG13 | 0.805 | 0.003 |
| I438 | N | 126.336 | 0.021 |
| I438 | QD1 | 0.372 | 0.011 |
| I438 | QG2 | 0.774 | 0.006 |
| T439 | C | 173.985 | 0.052 |
| T439 | CA | 62.111 | 0.056 |
| T439 | CB | 65.530 | 0.026 |
| T439 | CG2 | 19.947 | 0.035 |
| T439 | H | 8.140 | 0.007 |
| T439 | HA | 3.753 | 0.005 |
| T439 | HB | 4.183 | 0.010 |
| T439 | N | 110.219 | 0.016 |
| T439 | QG2 | 1.317 | 0.002 |
| L440 | C | 176.094 | 0.019 |
| L440 | CA | 54.846 | 0.061 |
| L440 | CB | 39.256 | 0.040 |
| L440 | CD1 | 24.176 | 0.061 |
| L440 | CD2 | 20.910 | 0.014 |
| L440 | CG | 24.588 | 0.000 |
| L440 | H | 7.111 | 0.005 |
| L440 | HA | 4.169 | 0.005 |
| L440 | HB2 | 1.884 | 0.005 |
| L440 | HB3 | 1.695 | 0.012 |
| L440 | HG | 1.427 | 0.005 |
| L440 | N | 122.127 | 0.019 |
| L440 | QD1 | 0.950 | 0.003 |
| L440 | QD2 | 1.008 | 0.004 |
| I441 | C | 174.010 | 0.031 |
| I441 | CA | 59.082 | 0.060 |
| I441 | CB | 34.362 | 0.059 |
| I441 | CD1 | 9.735 | 0.008 |
| I441 | CG1 | 26.091 | 0.023 |
| I441 | CG2 | 16.167 | 0.041 |
| I441 | H | 6.645 | 0.009 |
| I441 | HA | 2.535 | 0.010 |
| I441 | HB | 1.851 | 0.009 |
| I441 | HG12 | 1.346 | 0.006 |
| I441 | HG13 | 1.289 | 0.009 |
| I441 | N | 118.862 | 0.047 |
| I441 | QD1 | 0.745 | 0.011 |

| Residue | Atom | Chemical Shift [ppm] | Standard Deviation [ppm] |
|---|---|---|---|
| I441 | QG2 | 0.890 | 0.004 |
| A442 | C | 174.334 | 0.019 |
| A442 | CA | 51.042 | 0.032 |
| A442 | CB | 15.935 | 0.029 |
| A442 | H | 8.176 | 0.004 |
| A442 | HA | 3.858 | 0.012 |
| A442 | N | 117.292 | 0.022 |
| A442 | QB | 1.256 | 0.006 |
| S443 | C | 172.377 | 0.023 |
| S443 | CA | 56.267 | 0.078 |
| S443 | CB | 61.957 | 0.055 |
| S443 | H | 7.129 | 0.005 |
| S443 | HA | 4.397 | 0.003 |
| S443 | HB2 | 4.079 | 0.001 |
| S443 | HB3 | 3.969 | 0.004 |
| S443 | N | 109.328 | 0.014 |
| F444 | C | 176.521 | 0.000 |
| F444 | CA | 52.665 | 0.075 |
| F444 | CB | 34.126 | 0.011 |
| F444 | CD1 | 128.372 | 0.022 |
| F444 | CE1 | 127.609 | 0.009 |
| F444 | CZ | 124.962 | 0.016 |
| F444 | H | 7.512 | 0.008 |
| F444 | HA | 4.809 | 0.007 |
| F444 | HB2 | 3.298 | 0.004 |
| F444 | HB3 | 3.472 | 0.009 |
| F444 | HZ | 6.785 | 0.002 |
| F444 | N | 123.645 | 0.044 |
| F444 | QD | 7.010 | 0.006 |
| F444 | QE | 6.777 | 0.005 |
| H445 | C | 175.135 | 0.000 |
| H445 | CA | 58.121 | 0.035 |
| H445 | CB | 27.971 | 0.010 |
| H445 | CD2 | 116.982 | 0.013 |
| H445 | CE1 | 136.233 | 0.010 |
| H445 | HA | 4.315 | 0.004 |
| H445 | HB2 | 3.182 | 0.005 |
| H445 | HB3 | 3.142 | 0.003 |
| H445 | HD2 | 7.026 | 0.008 |
| H445 | HE1 | 7.722 | 0.006 |
| R446 | C | 174.814 | 0.033 |
| R446 | CA | 55.985 | 0.045 |
| R446 | CB | 27.382 | 0.086 |
| R446 | CD | 40.253 | 0.042 |
| R446 | CG | 25.461 | 0.061 |
| R446 | H | 8.339 | 0.016 |
| R446 | HA | 3.942 | 0.008 |
| R446 | HG2 | 1.646 | 0.003 |
| R446 | HG3 | 1.536 | 0.009 |
| R446 | N | 114.474 | 0.011 |
| R446 | QB | 1.805 | 0.006 |
| R446 | QD | 3.080 | 0.003 |
| V447 | C | 174.638 | 0.046 |
| V447 | CA | 63.216 | 0.038 |
| V447 | CB | 28.945 | 0.046 |
| V447 | CG1 | 20.825 | 0.039 |
| V447 | CG2 | 20.121 | 0.091 |
| V447 | H | 7.343 | 0.011 |

| Residue | Atom | Chemical Shift [ppm] | Standard Deviation [ppm] |
|---|---|---|---|
| V447 | HA | 3.823 | 0.012 |
| V447 | HB | 2.463 | 0.012 |
| V447 | N | 116.796 | 0.036 |
| V447 | QG1 | 1.129 | 0.004 |
| V447 | QG2 | 0.966 | 0.007 |
| Q448 | C | 174.072 | 0.032 |
| Q448 | CA | 55.780 | 0.074 |
| Q448 | CB | 26.120 | 0.032 |
| Q448 | CD | 177.601 | 0.004 |
| Q448 | CG | 31.928 | 0.046 |
| Q448 | H | 8.412 | 0.009 |
| Q448 | HA | 4.240 | 0.014 |
| Q448 | HB2 | 2.133 | 0.003 |
| Q448 | HB3 | 1.986 | 0.006 |
| Q448 | HE21 | 7.482 | 0.009 |
| Q448 | HE22 | 6.811 | 0.006 |
| Q448 | HG2 | 2.479 | 0.002 |
| Q448 | HG3 | 2.338 | 0.002 |
| Q448 | N | 119.805 | 0.029 |
| Q448 | NE2 | 111.894 | 0.009 |
| A449 | C | 175.723 | 0.003 |
| A449 | CA | 51.155 | 0.039 |
| A449 | CB | 15.946 | 0.033 |
| A449 | H | 7.500 | 0.005 |
| A449 | HA | 4.002 | 0.005 |
| A449 | N | 116.670 | 0.022 |
| A449 | QB | 1.307 | 0.003 |
| L450 | C | 173.958 | 0.010 |
| L450 | CA | 53.361 | 0.076 |
| L450 | CB | 40.586 | 0.027 |
| L450 | CD1 | 23.686 | 0.009 |
| L450 | CD2 | 20.118 | 0.023 |
| L450 | CG | 24.077 | 0.005 |
| L450 | H | 7.626 | 0.004 |
| L450 | HA | 4.320 | 0.007 |
| L450 | HB2 | 2.155 | 0.006 |
| L450 | HB3 | 1.306 | 0.004 |
| L450 | HG | 1.857 | 0.005 |
| L450 | N | 116.825 | 0.025 |
| L450 | QD1 | 0.890 | 0.009 |
| L450 | QD2 | 0.821 | 0.001 |
| T451 | C | 168.305 | 0.001 |
| T451 | CA | 59.398 | 0.046 |
| T451 | CB | 65.640 | 0.029 |
| T451 | CG2 | 16.436 | 0.030 |
| T451 | H | 8.138 | 0.004 |
| T451 | HA | 4.587 | 0.013 |
| T451 | HB | 4.269 | 0.004 |
| T451 | N | 111.223 | 0.016 |
| T451 | QG2 | 0.900 | 0.003 |
| T452 | C | 171.346 | 0.010 |
| T452 | CA | 58.025 | 0.064 |
| T452 | CB | 66.292 | 0.069 |
| T452 | CG2 | 18.881 | 0.021 |
| T452 | H | 8.077 | 0.006 |
| T452 | HA | 4.496 | 0.003 |
| T452 | HB | 4.612 | 0.011 |
| T452 | N | 113.504 | 0.019 |

| Residue | Atom | Chemical Shift [ppm] | Standard Deviation [ppm] |
|---|---|---|---|
| T452 | QG2 | 1.236 | 0.005 |
| D453 | C | 173.631 | 0.006 |
| D453 | CA | 50.325 | 0.051 |
| D453 | CB | 38.315 | 0.033 |
| D453 | H | 8.938 | 0.006 |
| D453 | HA | 4.712 | 0.008 |
| D453 | HB2 | 3.117 | 0.006 |
| D453 | HB3 | 2.505 | 0.009 |
| D453 | N | 126.062 | 0.019 |
| I454 | C | 174.588 | 0.027 |
| I454 | CA | 60.595 | 0.075 |
| I454 | CB | 34.878 | 0.071 |
| I454 | CD1 | 10.018 | 0.056 |
| I454 | CG1 | 26.111 | 0.033 |
| I454 | CG2 | 15.290 | 0.022 |
| I454 | H | 8.581 | 0.008 |
| I454 | HA | 3.727 | 0.008 |
| I454 | HB | 1.926 | 0.011 |
| I454 | HG12 | 1.445 | 0.002 |
| I454 | HG13 | 1.351 | 0.009 |
| I454 | N | 126.920 | 0.018 |
| I454 | QD1 | 0.736 | 0.014 |
| I454 | QG2 | 0.914 | 0.005 |
| S455 | C | 174.841 | 0.060 |
| S455 | CA | 59.482 | 0.049 |
| S455 | CB | 59.739 | 0.022 |
| S455 | H | 8.292 | 0.006 |
| S455 | HA | 4.362 | 0.006 |
| S455 | N | 117.270 | 0.019 |
| S455 | QB | 4.006 | 0.005 |
| L456 | C | 175.820 | 0.012 |
| L456 | CA | 54.392 | 0.027 |
| L456 | CB | 39.201 | 0.036 |
| L456 | CD1 | 22.505 | 0.025 |
| L456 | CD2 | 21.624 | 0.026 |
| L456 | CG | 24.698 | 0.014 |
| L456 | H | 7.484 | 0.007 |
| L456 | HA | 4.256 | 0.010 |
| L456 | HB2 | 1.898 | 0.011 |
| L456 | HB3 | 1.633 | 0.015 |
| L456 | HG | 1.579 | 0.002 |
| L456 | N | 126.114 | 0.018 |
| L456 | QD1 | 0.906 | 0.007 |
| L456 | QD2 | 0.944 | 0.004 |
| I457 | C | 174.916 | 0.008 |
| I457 | CA | 63.475 | 0.085 |
| I457 | CB | 34.240 | 0.096 |
| I457 | CD1 | 10.922 | 0.020 |
| I457 | CG1 | 26.552 | 0.016 |
| I457 | CG2 | 14.278 | 0.049 |
| I457 | H | 7.428 | 0.007 |
| I457 | HA | 3.503 | 0.009 |
| I457 | HB | 1.966 | 0.009 |
| I457 | HG12 | 1.718 | 0.001 |
| I457 | HG13 | 0.684 | 0.015 |
| I457 | N | 119.683 | 0.017 |
| I457 | QD1 | 0.679 | 0.004 |
| I457 | QG2 | 0.568 | 0.001 |

| Residue | Atom | Chemical Shift [ppm] | Standard Deviation [ppm] |
|---|---|---|---|
| F458 | C | 176.411 | 0.011 |
| F458 | CA | 59.539 | 0.055 |
| F458 | CB | 36.234 | 0.037 |
| F458 | CD1 | 128.830 | 0.006 |
| F458 | CE1 | 128.873 | 0.012 |
| F458 | CZ | 126.955 | 0.013 |
| F458 | H | 8.035 | 0.005 |
| F458 | HA | 4.002 | 0.006 |
| F458 | HZ | 7.183 | 0.003 |
| F458 | N | 116.234 | 0.017 |
| F458 | QB | 3.104 | 0.007 |
| F458 | QD | 7.306 | 0.004 |
| F458 | QE | 7.294 | 0.003 |
| A459 | C | 177.869 | 0.009 |
| A459 | CA | 52.518 | 0.065 |
| A459 | CB | 15.777 | 0.015 |
| A459 | H | 7.859 | 0.005 |
| A459 | HA | 4.108 | 0.004 |
| A459 | N | 119.891 | 0.023 |
| A459 | QB | 1.567 | 0.003 |
| A460 | C | 175.944 | 0.005 |
| A460 | CA | 51.830 | 0.074 |
| A460 | CB | 16.726 | 0.027 |
| A460 | H | 8.519 | 0.010 |
| A460 | HA | 4.033 | 0.011 |
| A460 | N | 119.964 | 0.021 |
| A460 | QB | 1.644 | 0.003 |
| L461 | C | 175.295 | 0.005 |
| L461 | CA | 51.707 | 0.060 |
| L461 | CB | 40.152 | 0.019 |
| L461 | CD1 | 23.076 | 0.017 |
| L461 | CD2 | 19.563 | 0.019 |
| L461 | CG | 23.470 | 0.003 |
| L461 | H | 7.433 | 0.003 |
| L461 | HA | 4.224 | 0.011 |
| L461 | HB2 | 1.775 | 0.011 |
| L461 | HB3 | 1.552 | 0.009 |
| L461 | HG | 1.991 | 0.003 |
| L461 | N | 112.371 | 0.022 |
| L461 | QD1 | 0.760 | 0.001 |
| L461 | QD2 | 0.817 | 0.002 |
| K462 | C | 173.792 | 0.029 |
| K462 | CA | 57.003 | 0.093 |
| K462 | CB | 29.927 | 0.090 |
| K462 | CD | 26.663 | 0.002 |
| K462 | CE | 39.453 | 0.000 |
| K462 | CG | 21.743 | 0.016 |
| K462 | H | 7.435 | 0.004 |
| K462 | HA | 3.959 | 0.011 |
| K462 | HB2 | 1.973 | 0.002 |
| K462 | HB3 | 1.825 | 0.003 |
| K462 | N | 121.660 | 0.018 |
| K462 | QD | 1.727 | 0.003 |
| K462 | QE | 3.029 | 0.010 |
| K462 | QG | 1.512 | 0.006 |
| D463 | C | 173.764 | 0.020 |
| D463 | CA | 50.749 | 0.068 |
| D463 | CB | 38.089 | 0.018 |

| Residue | Atom | Chemical Shift [ppm] | Standard Deviation [ppm] |
|---|---|---|---|
| D463 | H | 8.175 | 0.005 |
| D463 | HA | 4.818 | 0.006 |
| D463 | HB2 | 2.767 | 0.006 |
| D463 | HB3 | 2.714 | 0.013 |
| D463 | N | 116.194 | 0.012 |
| S464 | C | 173.731 | 0.006 |
| S464 | CA | 56.424 | 0.028 |
| S464 | CB | 61.507 | 0.029 |
| S464 | H | 7.563 | 0.006 |
| S464 | HA | 4.266 | 0.002 |
| S464 | HB2 | 4.137 | 0.013 |
| S464 | HB3 | 3.864 | 0.007 |
| S464 | N | 113.075 | 0.013 |
| K465 | C | 173.923 | 0.003 |
| K465 | CA | 53.731 | 0.021 |
| K465 | CB | 29.882 | 0.030 |
| K465 | CD | 26.035 | 0.024 |
| K465 | CE | 39.636 | 0.015 |
| K465 | CG | 22.653 | 0.015 |
| K465 | H | 9.600 | 0.010 |
| K465 | HA | 4.387 | 0.010 |
| K465 | HB2 | 2.064 | 0.003 |
| K465 | HB3 | 1.772 | 0.004 |
| K465 | HG2 | 1.590 | 0.002 |
| K465 | HG3 | 1.521 | 0.003 |
| K465 | N | 127.073 | 0.016 |
| K465 | QD | 1.703 | 0.003 |
| K465 | QE | 3.045 | 0.004 |
| V466 | C | 173.887 | 0.006 |
| V466 | CA | 61.794 | 0.031 |
| V466 | CB | 32.377 | 0.008 |
| V466 | CG1 | 18.921 | 0.072 |
| V466 | CG2 | 18.190 | 0.000 |
| V466 | H | 8.369 | 0.004 |
| V466 | HA | 3.961 | 0.004 |
| V466 | HB | 1.737 | 0.009 |
| V466 | N | 119.446 | 0.029 |
| V466 | QQG | 0.956 | 0.004 |
| V467 | C | 170.961 | 0.016 |
| V467 | CA | 55.925 | 0.058 |
| V467 | CB | 31.842 | 0.033 |
| V467 | CG1 | 19.381 | 0.032 |
| V467 | CG2 | 16.096 | 0.021 |
| V467 | H | 7.770 | 0.004 |
| V467 | HA | 4.888 | 0.010 |
| V467 | HB | 1.724 | 0.004 |
| V467 | N | 112.316 | 0.030 |
| V467 | QG1 | 0.241 | 0.008 |
| V467 | QG2 | 0.661 | 0.004 |
| E468 | C | 170.951 | 0.008 |
| E468 | CA | 51.218 | 0.045 |
| E468 | CB | 30.729 | 0.066 |
| E468 | CG | 32.583 | 0.018 |
| E468 | H | 8.963 | 0.009 |
| E468 | HA | 4.708 | 0.007 |
| E468 | HG2 | 2.091 | 0.002 |
| E468 | HG3 | 1.946 | 0.011 |
| E468 | N | 120.874 | 0.027 |

| Residue | Atom | Chemical Shift [ppm] | Standard Deviation [ppm] |
|---|---|---|---|
| E468 | QB | 1.735 | 0.006 |
| I469 | C | 173.944 | 0.031 |
| I469 | CA | 56.779 | 0.067 |
| I469 | CB | 37.108 | 0.031 |
| I469 | CD1 | 11.791 | 0.006 |
| I469 | CG1 | 24.672 | 0.048 |
| I469 | CG2 | 14.916 | 0.059 |
| I469 | H | 8.506 | 0.006 |
| I469 | HA | 5.158 | 0.009 |
| I469 | HB | 1.712 | 0.002 |
| I469 | HG12 | 1.543 | 0.001 |
| I469 | HG13 | 0.877 | 0.001 |
| I469 | N | 123.186 | 0.041 |
| I469 | QD1 | 0.785 | 0.008 |
| I469 | QG2 | 0.860 | 0.013 |
| V470 | C | 172.424 | 0.016 |
| V470 | CA | 59.279 | 0.059 |
| V470 | CB | 31.857 | 0.037 |
| V470 | CG1 | 18.412 | 0.000 |
| V470 | CG2 | 17.926 | 0.014 |
| V470 | H | 8.699 | 0.006 |
| V470 | HA | 4.085 | 0.008 |
| V470 | HB | 1.675 | 0.007 |
| V470 | N | 128.683 | 0.019 |
| V470 | QG1 | 0.947 | 0.010 |
| V470 | QG2 | 0.770 | 0.009 |
| D471 | C | 171.995 | 0.000 |
| D471 | CA | 53.862 | 0.053 |
| D471 | CB | 36.683 | 0.014 |
| D471 | H | 9.080 | 0.009 |
| D471 | HA | 4.100 | 0.005 |
| D471 | HB2 | 2.742 | 0.011 |
| D471 | HB3 | 2.665 | 0.002 |
| D471 | N | 127.084 | 0.033 |
| E472 | C | 172.130 | 0.000 |
| E472 | CA | 53.907 | 0.043 |
| E472 | CB | 25.777 | 0.029 |
| E472 | CG | 34.215 | 0.036 |
| E472 | H | 8.464 | 0.005 |
| E472 | HA | 4.359 | 0.013 |
| E472 | HB2 | 2.409 | 0.006 |
| E472 | HB3 | 2.197 | 0.008 |
| E472 | HG2 | 2.262 | 0.002 |
| E472 | HG3 | 2.174 | 0.003 |
| E472 | N | 116.407 | 0.000 |
| K473 | C | 171.170 | 0.027 |
| K473 | CA | 52.502 | 0.061 |
| K473 | CB | 32.934 | 0.032 |
| K473 | CD | 26.343 | 0.027 |
| K473 | CE | 38.883 | 0.044 |
| K473 | CG | 22.029 | 0.025 |
| K473 | H | 8.317 | 0.008 |
| K473 | HA | 5.131 | 0.012 |
| K473 | HB2 | 1.601 | 0.002 |
| K473 | HB3 | 1.548 | 0.001 |
| K473 | HE2 | 2.431 | 0.003 |
| K473 | HE3 | 2.086 | 0.004 |
| K473 | HG2 | 1.274 | 0.002 |

| Residue | Atom | Chemical Shift [ppm] | Standard Deviation [ppm] |
|---|---|---|---|
| K473 | HG3 | 1.213 | 0.005 |
| K473 | N | 121.131 | 0.023 |
| K473 | QD | 1.152 | 0.007 |
| K473 | QG | 1.239 | 0.014 |
| V474 | C | 170.956 | 0.050 |
| V474 | CA | 56.047 | 0.043 |
| V474 | CB | 32.992 | 0.064 |
| V474 | CG1 | 18.276 | 0.017 |
| V474 | CG2 | 18.988 | 0.031 |
| V474 | H | 9.366 | 0.006 |
| V474 | HA | 5.568 | 0.011 |
| V474 | HB | 1.722 | 0.007 |
| V474 | N | 117.386 | 0.024 |
| V474 | QG1 | 0.750 | 0.004 |
| V474 | QG2 | 0.631 | 0.007 |
| R475 | C | 171.298 | 0.033 |
| R475 | CA | 51.090 | 0.041 |
| R475 | CB | 32.364 | 0.058 |
| R475 | CD | 40.548 | 0.059 |
| R475 | CG | 20.849 | 0.011 |
| R475 | CZ | 156.147 | 0.000 |
| R475 | H | 8.428 | 0.013 |
| R475 | HA | 4.732 | 0.008 |
| R475 | HB2 | 1.454 | 0.005 |
| R475 | HB3 | 0.972 | 0.009 |
| R475 | HD2 | 2.366 | 0.005 |
| R475 | HD3 | 0.296 | 0.006 |
| R475 | HE | 7.596 | 0.007 |
| R475 | HG2 | 1.145 | 0.003 |
| R475 | HG3 | 0.888 | 0.005 |
| R475 | N | 119.583 | 0.055 |
| R475 | NE | 86.388 | 0.030 |
| R476 | C | 172.312 | 0.011 |
| R476 | CA | 54.134 | 0.059 |
| R476 | CB | 28.737 | 0.037 |
| R476 | CD | 41.025 | 0.022 |
| R476 | CG | 26.268 | 0.066 |
| R476 | CZ | 156.894 | 0.000 |
| R476 | H | 8.768 | 0.005 |
| R476 | HA | 4.126 | 0.007 |
| R476 | HB2 | 1.818 | 0.006 |
| R476 | HB3 | 1.694 | 0.003 |
| R476 | HD2 | 3.334 | 0.012 |
| R476 | HD3 | 2.985 | 0.006 |
| R476 | HE | 7.606 | 0.003 |
| R476 | N | 121.447 | 0.018 |
| R476 | NE | 84.254 | 0.018 |
| R476 | QG | 1.910 | 0.002 |
| R477 | C | 174.579 | 0.004 |
| R477 | CA | 56.839 | 0.062 |
| R477 | CB | 27.814 | 0.025 |
| R477 | CD | 40.339 | 0.053 |
| R477 | CG | 26.618 | 0.043 |
| R477 | CZ | 156.730 | 0.000 |
| R477 | H | 8.186 | 0.004 |
| R477 | HA | 3.783 | 0.010 |
| R477 | HB2 | 1.707 | 0.007 |
| R477 | HB3 | 1.401 | 0.003 |

| Residue | Atom | Chemical Shift [ppm] | Standard Deviation [ppm] |
|---|---|---|---|
| R477 | HD2 | 3.187 | 0.003 |
| R477 | HD3 | 2.912 | 0.004 |
| R477 | HE | 6.543 | 0.006 |
| R477 | HG2 | 1.415 | 0.009 |
| R477 | HG3 | 1.191 | 0.008 |
| R477 | N | 121.666 | 0.031 |
| R477 | NE | 83.298 | 0.019 |
| E478 | C | 171.995 | 0.011 |
| E478 | CA | 53.284 | 0.067 |
| E478 | CB | 27.914 | 0.027 |
| E478 | CG | 34.027 | 0.033 |
| E478 | H | 8.359 | 0.009 |
| E478 | HA | 4.334 | 0.002 |
| E478 | HB2 | 2.061 | 0.008 |
| E478 | HB3 | 1.932 | 0.004 |
| E478 | HG2 | 2.298 | 0.012 |
| E478 | HG3 | 2.137 | 0.001 |
| E478 | N | 118.729 | 0.030 |
| E479 | C | 170.615 | 0.000 |
| E479 | CA | 53.732 | 0.078 |
| E479 | CB | 27.254 | 0.035 |
| E479 | CG | 34.520 | 0.029 |
| E479 | H | 9.063 | 0.006 |
| E479 | HA | 4.079 | 0.005 |
| E479 | HB2 | 2.132 | 0.007 |
| E479 | HB3 | 2.049 | 0.001 |
| E479 | HG2 | 2.318 | 0.002 |
| E479 | HG3 | 2.224 | 0.002 |
| E479 | N | 123.103 | 0.019 |
| P480 | C | 176.963 | 0.009 |
| P480 | CA | 62.697 | 0.036 |
| P480 | CB | 28.664 | 0.059 |
| P480 | CD | 48.391 | 0.015 |
| P480 | CG | 24.465 | 0.000 |
| P480 | HA | 4.240 | 0.004 |
| P480 | HB2 | 2.947 | 0.007 |
| P480 | HB3 | 2.102 | 0.007 |
| P480 | HD2 | 3.738 | 0.009 |
| P480 | HD3 | 3.199 | 0.006 |
| P480 | QG | 1.861 | 0.009 |
| E481 | C | 174.214 | 0.023 |
| E481 | CA | 55.857 | 0.048 |
| E481 | CB | 25.320 | 0.040 |
| E481 | CG | 33.180 | 0.061 |
| E481 | H | 9.544 | 0.005 |
| E481 | HA | 3.817 | 0.006 |
| E481 | HB2 | 2.140 | 0.007 |
| E481 | HB3 | 2.022 | 0.002 |
| E481 | N | 114.869 | 0.012 |
| E481 | QG | 2.288 | 0.003 |
| K482 | C | 174.910 | 0.007 |
| K482 | CA | 54.869 | 0.065 |
| K482 | CB | 29.502 | 0.040 |
| K482 | CD | 26.384 | 0.031 |
| K482 | CE | 39.067 | 0.032 |
| K482 | CG | 21.207 | 0.035 |
| K482 | H | 7.410 | 0.007 |
| K482 | HA | 4.030 | 0.007 |

| Residue | Atom | Chemical Shift [ppm] | Standard Deviation [ppm] |
|---|---|---|---|
| K482 | HB2 | 1.312 | 0.004 |
| K482 | HB3 | 1.046 | 0.006 |
| K482 | HE2 | 2.482 | 0.004 |
| K482 | HE3 | 2.445 | 0.014 |
| K482 | HG2 | 0.757 | 0.009 |
| K482 | HG3 | 0.479 | 0.008 |
| K482 | N | 118.914 | 0.028 |
| K482 | QD | 1.283 | 0.014 |
| W483 | C | 169.919 | 0.000 |
| W483 | CA | 56.223 | 0.040 |
| W483 | CB | 28.537 | 0.051 |
| W483 | CD1 | 125.360 | 0.024 |
| W483 | CE2 | 137.240 | 0.000 |
| W483 | CE3 | 116.820 | 0.019 |
| W483 | CH2 | 122.495 | 0.017 |
| W483 | CZ2 | 112.240 | 0.021 |
| W483 | CZ3 | 119.191 | 0.011 |
| W483 | H | 6.854 | 0.006 |
| W483 | HA | 4.609 | 0.006 |
| W483 | HB2 | 3.348 | 0.006 |
| W483 | HB3 | 3.274 | 0.010 |
| W483 | HD1 | 7.471 | 0.006 |
| W483 | HE1 | 10.669 | 0.008 |
| W483 | HE3 | 7.164 | 0.005 |
| W483 | HH2 | 7.268 | 0.005 |
| W483 | HZ2 | 7.487 | 0.005 |
| W483 | HZ3 | 6.847 | 0.008 |
| W483 | N | 120.774 | 0.034 |
| W483 | NE1 | 129.872 | 0.019 |
| P484 | C | 173.792 | 0.002 |
| P484 | CA | 62.947 | 0.038 |
| P484 | CB | 28.688 | 0.075 |
| P484 | CD | 47.829 | 0.017 |
| P484 | CG | 26.991 | 0.034 |
| P484 | HA | 4.628 | 0.009 |
| P484 | HB2 | 2.235 | 0.003 |
| P484 | HB3 | 1.659 | 0.009 |
| P484 | HD2 | 4.129 | 0.002 |
| P484 | HD3 | 3.518 | 0.010 |
| P484 | HG2 | 2.188 | 0.009 |
| P484 | HG3 | 1.963 | 0.005 |
| L485 | C | 170.753 | 0.000 |
| L485 | CA | 48.535 | 0.038 |
| L485 | CB | 42.872 | 0.032 |
| L485 | CD1 | 23.487 | 0.030 |
| L485 | CD2 | 20.972 | 0.019 |
| L485 | CG | 24.183 | 0.000 |
| L485 | H | 7.573 | 0.005 |
| L485 | HA | 4.978 | 0.008 |
| L485 | HB2 | 1.604 | 0.013 |
| L485 | HB3 | 1.316 | 0.004 |
| L485 | HG | 1.649 | 0.013 |
| L485 | N | 125.904 | 0.014 |
| L485 | QD1 | 0.912 | 0.004 |
| L485 | QD2 | 0.983 | 0.003 |
| P486 | CA | 61.671 | 0.017 |
| P486 | CB | 29.508 | 0.016 |
| P486 | CD | 47.532 | 0.007 |

| Residue | Atom | Chemical Shift [ppm] | Standard Deviation [ppm] |
|---|---|---|---|
| P486 | CG | 24.296 | 0.000 |
| P486 | HA | 4.346 | 0.002 |
| P486 | HB2 | 2.223 | 0.009 |
| P486 | HB3 | 1.952 | 0.009 |
| P486 | QD | 3.656 | 0.003 |
| P486 | QG | 1.980 | 0.005 |
